# CFIm25 regulates human stem cell function independently of its role in mRNA alternative polyadenylation

**DOI:** 10.1101/2021.12.08.471721

**Authors:** Yi Ran, Shanshan Huang, Junjie Shi, Qiumin Feng, Yanhui Deng, Andy Peng Xiang, Chengguo Yao

## Abstract

It has recently been shown that CFIm25, a canonical mRNA 3’ processing factor, could play a variety of physiological roles through its molecular function in the regulation of mRNA alternative polyadenylation (APA). Here, we used CRISPR/Cas9-mediated gene editing approach in human embryonic stem cells (hESCs) for CFIm25, and obtained three gene knockdown/mutant cell lines. CFIm25 gene editing resulted in higher proliferation rate and impaired differentiation potential for hESCs, with these effects likely to be directly regulated by the target genes, including the pluripotency factor *rex1*. Mechanistically, we unexpected found that perturbation in CFIm25 gene expression did not significantly affect cellular mRNA 3’ processing efficiency and APA profile. Rather, we provided evidences that CFIm25 may impact RNA polymerase II (RNAPII) occupancy at the body of transcribed genes, and promote the expression level of a group of transcripts associated with cellular proliferation and/or differentiation. Further study indicated that CFIm25 association with LEO1, an RNAPII associated factor, might contribute to the effect. Taken together, these results reveal novel mechanisms underlying CFIm25’s modulation in determination of cell fate, and provide evidence that the process of mammalian gene transcription may be regulated by an mRNA 3’ processing factor.

## Introduction

Processing of pre-mRNA 3’ end is a key step in eukaryotic gene expression (Colgan & Manley, 1997). Based on current models, processing of human canonical mRNA 3’ end involves two coupled steps, namely cleavage and polyadenylation (Shi *et al*, 2009; Sun *et al*, 2020; Sun *et al*, 2018; Zhang *et al*, 2020). Specifically, cleavage requires two core multi-subunit complexes, namely cleavage and polyadenylation specificity factors (CPSF) and cleavage stimulation factor (CstF). On the other hand, polyadenylation involves addition of a poly(A) tail at the 3’ end of pre-mRNA upon cleavage by Poly(A) polymerase (PAP). At the molecular level, mRNA 3’ processing often occurs co-transcriptionally (Bentley, 2014; Fusby *et al*, 2016; Glover-Cutter *et al*, 2008), and is tightly connected with all the three steps of mRNA transcription, namely initiation, elongation, and termination. For example, TFIID, one of the general transcription factors required for transcription initiation, has been implicated in regulation of mRNA 3’ processing by associating with CPSF (Dantonel *et al*, 1997), the core subunit of 3’ processing complex. More recent studies have shown that transcription activity at 5’ end of genes could significantly affect mRNA 3’ processing, though the detailed mechanisms remain elusive (Ji *et al*, 2011; Nagaike *et al*, 2011; Rosonina *et al*, 2003). Another example is the phosphorylation of serine 2 residues (Ser2P) at the C-terminal domain (CTD) of heptad repeats of RPB1, the largest subunit of RNAPII. As transcription approaches termination, Ser2P facilitates recruitment of 3’ processing factors to nascent transcripts (Davidson *et al*, 2014; Licatalosi *et al*, 2002). Aside from the impact of transcription on mRNA 3’ processing, emerging evidences have shown that mRNA 3’ processing, in turn, might impact transcription. For example, several yeast 3’ processing factors reportedly interact with 5’ end of genes, thereby impacting transcription through gene looping (Al Husini *et al*, 2013; Allepuz-Fuster *et al*, 2019; Ansari & Hampsey, 2005; El Kaderi *et al*, 2009). In human cells, 3’ end formation has been shown to play a stimulatory role in transcription, possibly by recycling factors required for initiation/elongation (Mapendano *et al*, 2010). Another example is U1 snRNP telescripting, a phenomenon linking premature transcription termination with mRNA 3’ processing at numerous intronic polyadenylation sites (PASs) (Kaida *et al*, 2010; Ran *et al*, 2021; Venters *et al*, 2019). However, despite the crucial role played by mRNA 3’ processing in mRNA maturation and function, its benefits to transcription is often underestimated and less studied (Cavallaro *et al*, 2021; Mapendano *et al*., 2010).

The human CFIm complex, which comprises CFIm68/CFIm59 and CFIm25, was initially identified as a basic subunit of canonical 3’ processing complex (Ruegsegger *et al*, 1996; Ruegsegger *et al*, 1998; Shi *et al*., 2009). Recent studies suggest that it serves as an activator of canonical mRNA 3’ processing and is a master regulator of alternative polyadenylation (APA) (Kim *et al*, 2010; Kubo *et al*, 2006; Martin *et al*, 2012; Masamha *et al*, 2014; Zhu *et al*, 2018). Accumulating evidences have indicated that this complex might play a role in gene transcription. Firstly, CFIm, together with CPSF and CstF, can be cross-linked with transcription initiation region for transcribed genes (Calvo & Manley, 2003; Garrido-Lecca *et al*, 2016; Glover-Cutter *et al*., 2008; Katahira *et al*, 2013). Secondly, researchers used RNAPII ChIP-seq to reveal transcription changes in a subset of genes following depletion of CFIm (Tellier *et al*, 2018, 2019). To date, nothing is known on whether the CFIm complex can directly regulate transcription of genes. This is, at least partially, due to a global APA shift upon CFIm depletion in previously reported cell systems (Alcott *et al*, 2020; Kim *et al*., 2010; Masamha *et al*., 2014; Sommerkamp *et al*, 2020; Tan *et al*, 2018; Weng *et al*, 2020; Zhu *et al*., 2018), and the effect of CFIm depletion on gene transcription may be neglected.

Recent studies have implicated CFIm25, the key component of the CFIm complex, in development of multiple cancer types and determination of cell fate (Brumbaugh *et al*, 2018; Chu *et al*, 2019; Jafari Najaf Abadi *et al*, 2019; Tan *et al*., 2018). Given the primary molecular function of CFIm25 in regulation of mRNA 3’ processing and APA, most of the reported CFIm25-associated cellular phenotypes have been attributed to its role in PAS choice of target genes thus far (Alcott *et al*., 2020; Brumbaugh *et al*., 2018; Chu *et al*., 2019; Gennarino *et al*, 2015; Huang *et al*, 2018; Jafari Najaf Abadi *et al*., 2019; Masamha *et al*., 2014; Sommerkamp *et al*., 2020; Tan *et al*., 2018; Weng *et al*., 2020; Weng *et al*, 2019; Zhou *et al*, 2020). In the present study, we used CFIm25 knockdown/mutant H9 cell lines, to elucidate an alternative underlying mechanism through which CFIm25 participates in gene regulatory network. Our results revealed that CFIm25 depletion/mutation has little effect on efficiency of cellular mRNA 3’ processing and global APA profile in H9 cell lines. Strikingly, disruption of CFIm25 gene expression significantly impacted RNAPII binding, at transcribed genes, and down-regulated transcription output of several key genes associated with the phenotype, including *rex1* gene. Overall, these results reveal a potential role played by CFIm25 in regulation of gene transcription.

## Results

### Generation of three CFIm25 knockdown/mutation cell lines in hESCs (human embryonic stem cells) using CRISPR/Cas9 technology

CFIm25 was recently shown to be a determinant factor of cell fate in mouse cells (Brumbaugh *et al*., 2018). To examine its role in human stem cells, we initially wished to perform CFIm25 gene knockout (KO) in H9 cells, a commonly used human embryonic stem cell line, by using the episomal vector-based CRISPR/Cas9 technology (Xie *et al*, 2017). T7 endonuclease I assays revealed that transfection of target gRNAs resulted in a relatively high mutation rate at the CFIm25 gene locus (Figure 1-figure supplement 1A; Figure 1 source-data file 1). After antibiotics selection, we picked more than 300 clones for western blot analysis (Figure 1-source data file 2), and results showed that at least three of them showed gene KO using a CFIm25 primary antibody (sc-81109, santa cruz) (Figure 1A; Figure 1-source data file 3), which recognizes the N-terminus of CFIm25 protein. We noted that our gRNAs were designed near the start codon (Figure 1-figure supplement 1B), it is possible that these clones may harbor mutation at CFIm25 N-terminus, which may not be targeted by this antibody. Indeed, using another antibody that can recognize the full length CFIm25 (10322-1-AP, Proteintech), we observed a faint band near 25 kDa for the selected clones (the knockdown efficiency reached approximately 90% for all the three clones) (Figure 1A; Figure 1-source data file 3). We further applied DNA sequencing and found that each clone at least one allele has been deleted a multiple of 3 nts (Figure 1-figure supplement 1B), which results in 12 to 17 amino acids N-terminus deletion proteins. These results are in line with the observation that the molecular weight of the band in mutant cells is slightly smaller than that in control cells using the CFIm25 antibody from Proteintech (Figure 1A; Figure 1-source data file 3). Consistent with previous reports (Masamha *et al*., 2014; So *et al*, 2019), we observed that CFIm59, but not CFIm68, showed a mild decrease in expression level upon CFIm25 depletion (Figure 1A; Figure 1-source data file 3). Taken together, we presumed that CFIm25 is essential for human cells, and we generated three CFIm25 gene knockdown and small N-terminus deletion mutant clones in H9 cells. For simplicity, they were designated as CFIm25-mutants (CFIm25 m). As mock controls in subsequent experiments, we randomly picked two clones that were transfected with CRISPR/Cas9 empty vector.

**Figure 1.**
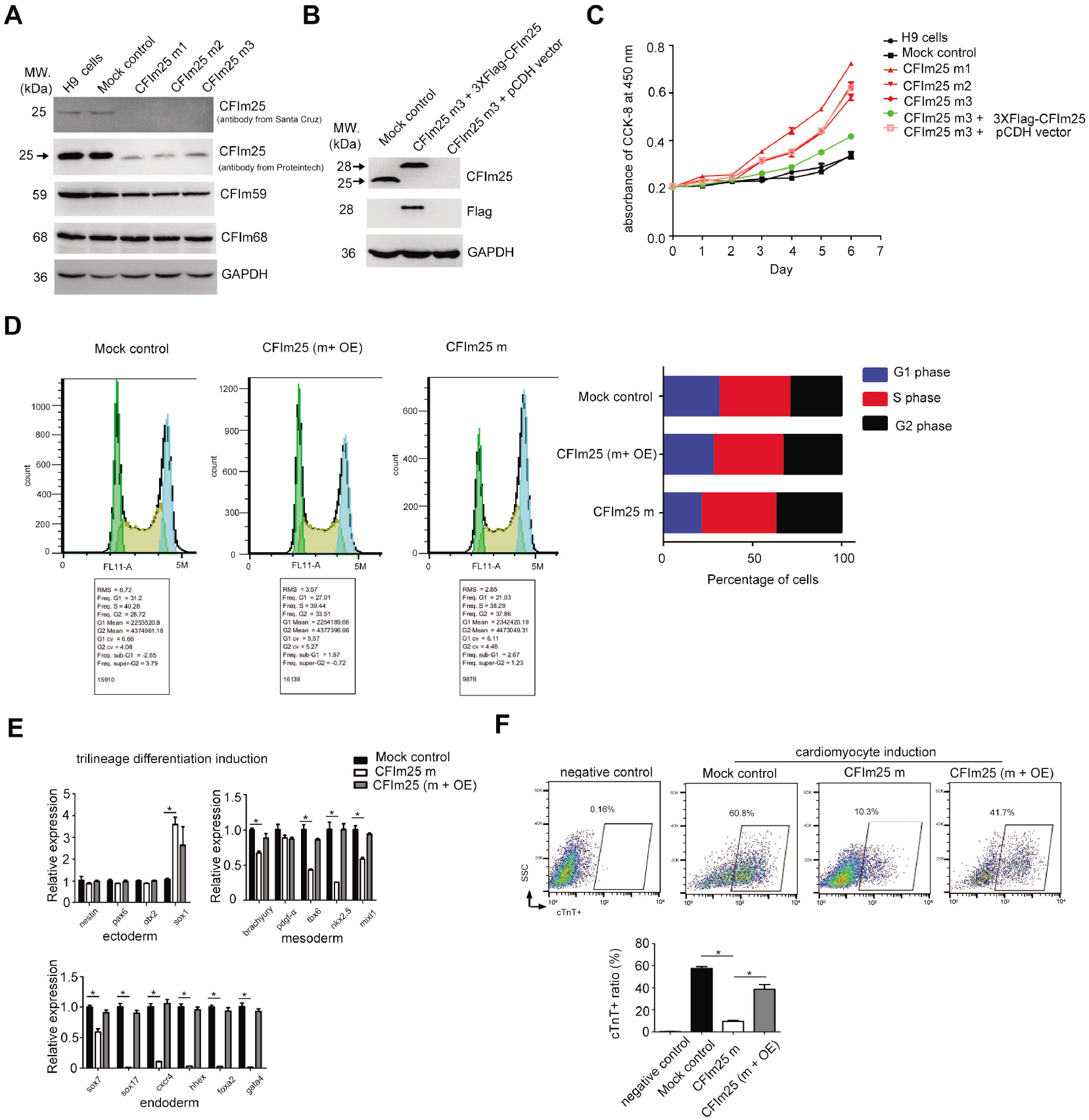
CFIm25 knockdown/mutation impacts the cell proliferation rate and differentiation potential of H9 cell line. (A) Western blot analysis of CFIm25 and CFIm59/68 proteins in cell lysates prepared from two controls and three CFIm25 gene-edited H9 cell lines. GAPDH serves as sample loading control (Control: H9 cells; Mock control: eCRISPR empty vector-transfected H9 cells; CFIm25 m1-3: eCRISPR-CFIm25 gRNAs transfected H9 cells). The primary antibody 1 against CFIm25 is from Santa Cruz Company, and antibody 2 is from Proteintech. (B) Western blot analysis of CFIm25 and indicated protein (peptide) in cell lysates prepared from Mock, CFIm25-m, and CFIm25-m plus 3XFIag-CFIm25 overexpression cells. At least three independent experiments have been carried out and representative images are shown. The primary antibody for CFIm25 is from santa cruz (sc-81109). (C) Cell proliferation rate measurement by CCK-8 kit for indicated cell lines. The growth rate of hESCs is largely dependent on starting cell density. The starting cell density in this experiment is 5000 per well of 96 well plates. Three independent experiments have been carried out and representative results are shown. (D) Flow Cytometry analysis of cell cycle using a Propidium Iodide Flow Cytometry Kit in the indicated cell lines (m: mutant; OE:overexpression). The right panel shows the representative result of the percentages of cells during different stages of cell cycle. The quantification of three independent experiments is shown in Figure 1-figure supplemental 1D. (E) RT-qPCR analysis of the expression level of corresponding lineage differentiation markers in indicated cell lines during trilineage differentiation. Three independent experiments have been carried out and quantified. Student’s t-test was used to estimate the significance of the change. *p<0.05. ns: non-significant. (F) Quantification of the yield of cardiomyocytes by performing fluorescence-activated cell sorting (FACS) analysis in the indicated cell lines during cardiomyocytes differentiation from three independent experiments. cTnT antibodies were used in FACS experiment. Bottom panel is the quantification from three representative experiments (m: mutant; OE: overexpression). Student’s t-test was used to estimate the significance: * p < 0.05.

### CFIm25 regulates growth rate and pluripotency of hESCs

During cell culture, we observed that all of these mutant cell lines grew faster than mock control cells. Therefore, we decided to decipher the potential physiological roles of CFIm25 in hESCs using these cells. Cell proliferation assessment using Cell Counting Kit-8 (CCK-8) showed that CFIm25 mutation caused a significant increase in cell proliferation rate in all three clones (Figure 1C). It is important to note that the growth rate of human stem cell is largely dependent on the starting cell density, we repeated the same experiment with another starting cell density, and the results showed similar trend (Figure 1-figure supplement 1C). This observation is consistent with previous reports that CFIm25 knock-down increased the rate of cell proliferation in multiple cancer cell lines (Jafari Najaf Abadi *et al*., 2019; Masamha *et al*., 2014), and suggests that ESCs could be sharing a common feature with cancer cells. Notably, since all three mutant clones were obtained following transfection of three independent gRNAs (Figure 1-figure supplement 1B), the observed phenotype may not be due to potential indirect effects caused by gRNA off-targeting. Consequently, we combined all three into one dataset (CFm25-m) for simplicity, it not indicated otherwise, owing to the high similarity among phenotypes and deep sequencing results (Figure 1C; Supplemental Table 2 and 3).

To elucidate the role of CFIm25 in proliferation of hESCs, we performed a rescue experiment by re-expressing CFIm25 in the mutant cells (CFIm25-m3) using a lentivirus-mediated gene overexpression system. Western blotting revealed that CFIm25 expression was restored to a level comparable to that of endogenous protein (Figure 1B; Figure 1-source data file 4), thereby re-establishing the cell proliferation phenotype (Figure 1C). Additionally, results from cell cycle analysis showed that mutation in CFIm25 significantly shortened the G1 phase and lengthened the G2 phase during cell cycle progression (Figure 1D; Figure 1-figure supplement 1D). Overall, these results demonstrated that CFIm25 plays an active role in proliferation of hESCs.

Next, we investigated whether CFIm25 might affect hESCs self-renewal capacity and differentiation potential, two main features characteristic of ESCs. To test this, we first performed qRT-PCR analysis targeting a panel of canonical pluripotency and differentiation markers in both mock control and CFIm25-mutant hESCs. Results revealed no significant changes in expression of the tested pluripotency markers (Figure 1-figure supplement 1E and 1F). This is consistent with the observation from cell morphology analysis (Figure 1-figure supplement 1G), and suggests that mutations in CFIm25 might not affect self-renewal capacity of hESCs. In keeping with this, the expression of marker genes across the three germ layers remained undetectable (Supplemental Table 1). Additionally, we used a well-defined Trilineage Differentiation Kit followed by qRT-PCR analysis of differentiation markers to compare the differentiation potential of mock control and CFIm25-mutant hESCs. Strikingly, mutation in CFIm25 appeared to interfere with endoderm, and to a lesser extent, mesoderm differentiation, as evidenced by downregulation of all tested endoderm markers as well as some in the mesoderm markers (Figure 1E).

To validate the positive role played by CFIm25 in hESC mesoderm/endoderm differentiation, we used a Cardiomyocyte Differentiation kit to generate cardiomyocytes from mock control and CFIm25-mutant hESCs, owing to the fact that cardiomyocyte specification requires both primitive endoderm and nascent mesoderm (Rowton *et al*, 2021; Ruan *et al*, 2019). Cardiomyocyte induction resulted in a ~5-fold decrease in efficiency of CFIm25-mutants, as evidenced by the percentages of cardiac troponin T-positive (cTnT+) cells (Figure 1F). Additionally, cardiomyocytes derived from mock control hESCs exhibited spontaneous beating on day 15, whereas less activity was observed in those from CFIm25-mutant hESCs (Supplemental Video 1 and 2). CFIm25 re-expression in CFIm25-mutant hESCs increased the induction efficiency by about 4 fold (Figure 1F), further affirming CFIm25’s role in cardiomyocytes. Taken together, these results indicated that CFIm25 regulates cell proliferation and differentiation potential in H9 cell line.

### CFIm25 regulates mRNA expression level in a subset of genes in hESCs

Next, we explored the molecular mechanisms through which CFIm25 regulates hESCs proliferation and pluripotency. Given its function in choice of polyadenylation site (PAS) and APA regulation (Brumbaugh *et al*., 2018; Jafari Najaf Abadi *et al*., 2019; Kubo *et al*., 2006), we hypothesized that the observed phenotype is, at least in part, caused by aberrant APA profile in CFIm25-mutant cells. To test this, we characterized global polyadenylation profiles in CFIm25-mutant hESCs alongside controls via high-throughput mRNA 3’ end sequencing. Unexpectedly, results showed that mutations in CFIm25 induced insignificant APA changes in the hESC transcriptome (Figure 2A; Supplemental Table 2), in contrast with what has previously been reported (Alcott *et al*., 2020; Brumbaugh *et al*., 2018; Chu *et al*., 2019; Huang *et al*., 2018; Masamha *et al*., 2014; Weng *et al*., 2019). As can be seen in Figure 2-figure supplement 1A, two well documented CFIm25 APA target, *cyclinD1* and *dicer1* genes, did not show apparent APA shift in CFIm25-mutant cells. Previous studies have shown that CFIm25-mediated APA regulation is associated with enhanced canonical PAS processing (Zhu *et al*., 2018). Therefore, we compared the overall canonical PAS processing efficiency between CFIm25-mutant and mock control hESCs using a luciferase reporter assay that is applied elsewhere (Lackford *et al*, 2014; Shi *et al*, 2019; Yao *et al*, 2012; Zhu *et al*., 2018). Results indicated that mutations in CFIm25 did not lower the processing activity of SVL PAS (Figure 2-figure supplement 1B), a widely used canonical PAS in the mRNA processing field. These results suggest that CFIm25-mutant is sufficient to support canonical mRNA 3’ processing in cells. To verify these findings, we performed a SVL PAS RNA-biotin based pull-down assay followed by western blot analysis using nuclear extracts (NEs) in CFIm25-mutant and mock control hESCs, and observed that core 3’ processing factors, such as CFIm68, Fip1 and CPSF30, were pulled down with similar efficiency (Figure 2-figure supplement 1C; Figure 2-source file 1). As negative controls, much less proteins were detected in the pull-down sample using SVL PAS RNA mutant, which harbors a point mutation at the core AAUAAA hexamer (Figure 2-figure supplement 1C). Consistently, the SVL PAS RNA mutant showed little PAS processing activity in vivo (Figure 2-figure supplement 1B).

**Figure 2.**
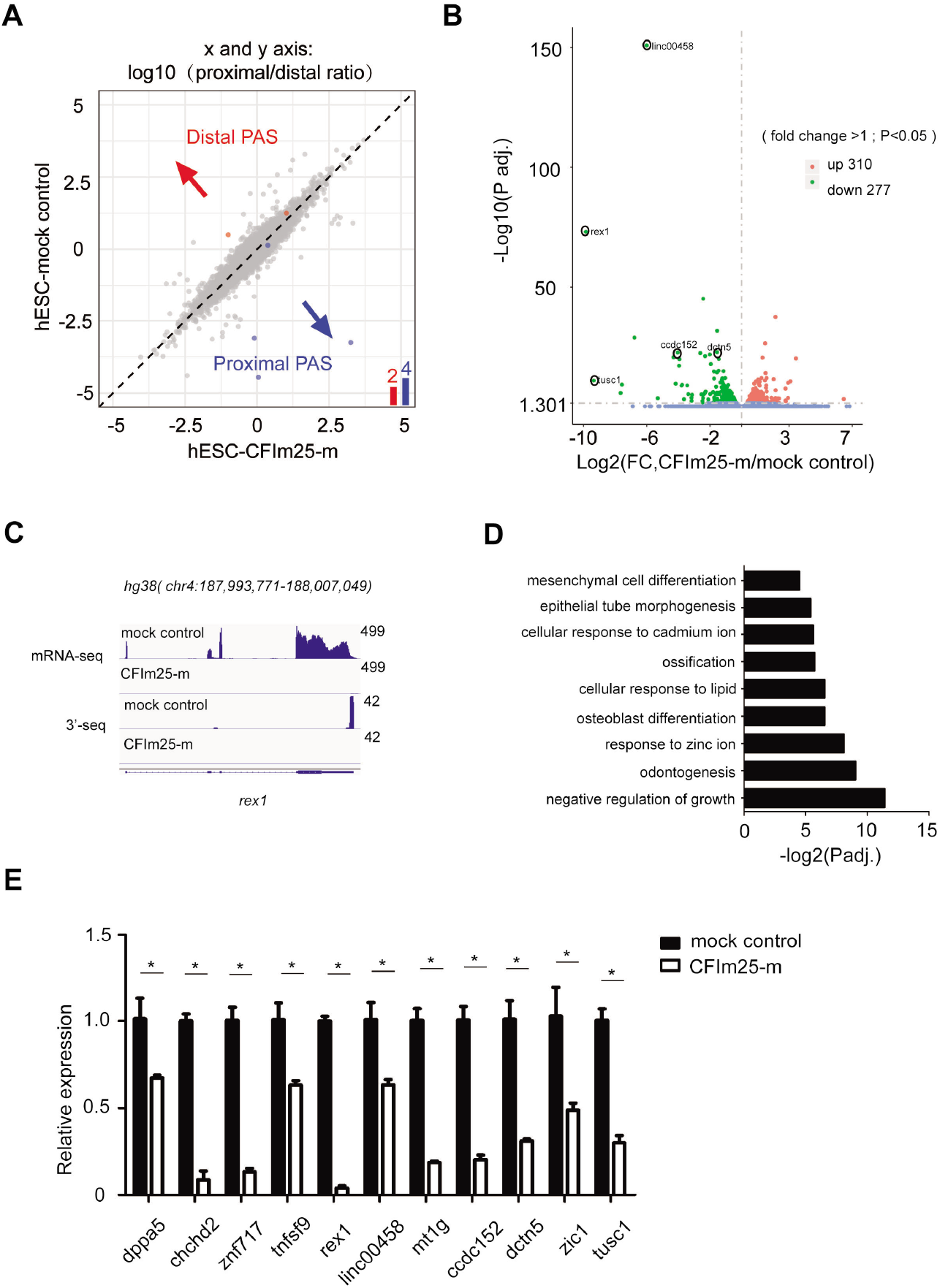
Effect of CFIm25 knockdown/mutation on the global mRNA alternative polyadenylation (APA) profile and expression level of poly(A+) transcripts in H9 cell line. (A) mRNA alternative polyadenylation (APA) change in mock and CFIm25-m(1-3) H9 cell lines. 3’-seq analysis of APA in mock and CFIm25-m H9 cells, Log2(proximal/distal ratio) are plotted for mock (y-axis) and CFIm25-m H9 cells (x-axis). Statistically significant changes are highlighted in blue (distal to proximal shift) and red (proximal to distal shift). The numbers of APA changes are shown in the column graph. (B) Volcano plot showing the expression level change of poly(A+) mRNA in mock and CFIm25-m (1-3) H9 cells. Significant changes (p<0.05, fold change>l) were colored red (up-regulated in CFIm25-m cells in comparison to mock cells) or green (down-regulated in CFIm25-m cells in comparison to mock cells), blue dots shows the changes either not statistically significant (p>0.05) or less reliable (fold change<1). Genes for subsequent studies are circled. (C) IGV track screen shots showing mRNA-seq and 3’-seq results for rex1 gene in mock and CFIm25-m H9 cells. (D) Gene ontology analysis of the group of down-regulated genes (277 genes) upon CFIm25-m using the Gene Ontology Consortium platform (http://geneontology.org/). Gene ontology terms (y axis) and corresponding p-values (x axis) are shown. (E) RT-qPCR analysis of the expression level of indicated genes in mock and CFIm25-m (1-3) H9 cells. The results of three independent experiments have been quantified. Student’s t-test was used to estimate the significance of the change. *P<0.05.

To further unravel the mechanisms underlying the cellular phenotype, we performed RNA-seq analysis to determine differential expression of genes between CFIm25-mutant and control H9 cells. At a cutoff value of P<0.05 and fold change>1, we found a total of 587 differentially expressed genes between the groups, of which 277 and 310 were down-regulated and up-regulated, respectively. On the other hand, 99 and 129 genes were down-regulated and up-regulated, respectively, at a cutoff value of P<0.05 and fold change>2 (Figure 2B; Supplemental Table 3). Additionally, both RNA-seq and PAS-seq approaches were efficient in quantification of gene/isoform expression, as evidenced by good agreement between respective results (Figure 2C; Figure 2-figure supplement 1D, 1E). Gene ontology analysis of the 99/277 down-regulated genes revealed significantly enrichment of genes involved in cellular differentiation, as well as development and negative regulation of growth (Figure 2D), which is consistent with the earlier results on phenotypes in CFIm25-mutant cells (Figure 1C-1F). In contrast, enrichment analysis of the 128/309 up-regulated genes revealed no terms associated with cellular proliferation or differentiation (Supplemental Table 3).

To further validate the RNA-seq results, we performed qRT-PCR analysis targeting 10 down-regulated genes, and found consistent expression patterns (Figure 2C, 2E; Figure 2-figure supplement 1E). Analysis of RNA-seq data from the aforementioned CFIm25 over-expression and mock control hESCs revealed that CFm25 re-expression restored expression of most down-regulated genes (Figure 2E; Figure 2-figure supplement 1F, 1G; Supplemental Table 4), suggesting that CFIm25 may be playing a direct role in regulating expression of these transcripts. Based on these findings, we hypothesized that CFIm25 might be regulating expression of a subset of cellular proliferation/differentiation-associated transcripts independent of its canonical role in promoting the processing of canonical PASs in hESCs.

### CFIm25 promotes *rex1* gene expression at the transcription level

Given that *rex1* is a well-established pluripotency marker and its expression showed the most significant change upon CFIm25 gene editing (Figure 2E) (Masui *et al*, 2008; Son *et al*, 2013), we further wished to understand how CFIm25 promotes *rex1* gene expression in hESCs. We considered several hypotheses. Firstly, *rex1* PAS 3’ processing might not be efficient in CFIm25-mutant cells, and may cause transcription read-through at the PAS region as well as subsequent mRNA decay; secondly, CFIm25 might protect *rex1* mRNA from degradation and promote its stability in the nucleus; and thirdly, CFIm25 may promote *rex1* gene transcription. To test the first hypothesis, we compared the levels of extended transcript beyond *rex1* PAS via qRT-PCR (Figure 3A), and observed that CFIm25 mutations did not increase the yield of read-through transcript at PAS region (Figure 3B). To test the second scenario, we measured the half-life of *rex1* mRNA by first treating cells with Actinyomycin D (Act D), followed by qRT-PCR analysis. Similarly, we found no marked difference between mock control and CFIm25-mutant cells within 2-hour periods (Figure 3-figure supplement 1A). In fact, *rex1* mRNA was hardly detectable in CFIm25-mutant cells following longer time Act D treatment. At least three lines of evidence suggest that CFIm25 regulates *rex1* gene expression at the transcriptional level. Firstly, qRT-PCR-based comparison of *rex1* pre-mRNA expression, using primers targeting its intronic region, revealed a fold change that was comparable to that of *rex1* mRNA expression (Figure 3C). Secondly, we enriched nascent RNAs by purifying chromatin-associated RNAs and subsequently quantified their expression via qRT-PCR using the aforementioned primers. We found consistent results, evidenced by lower levels of *rex1* pre-mRNAs in CFIm25-mutant, relative to control cells (Figure 3-figure supplement 1B). Thirdly, we directly enriched nascent RNAs via metabolic pulse-chase labeling of RNA using bromouridine (BrU), then purified them with anti-BrU antibody. As expected, we obtained similar results after qRT-PCR (Figure 3-figure supplement 1C).

**Figure 3.**
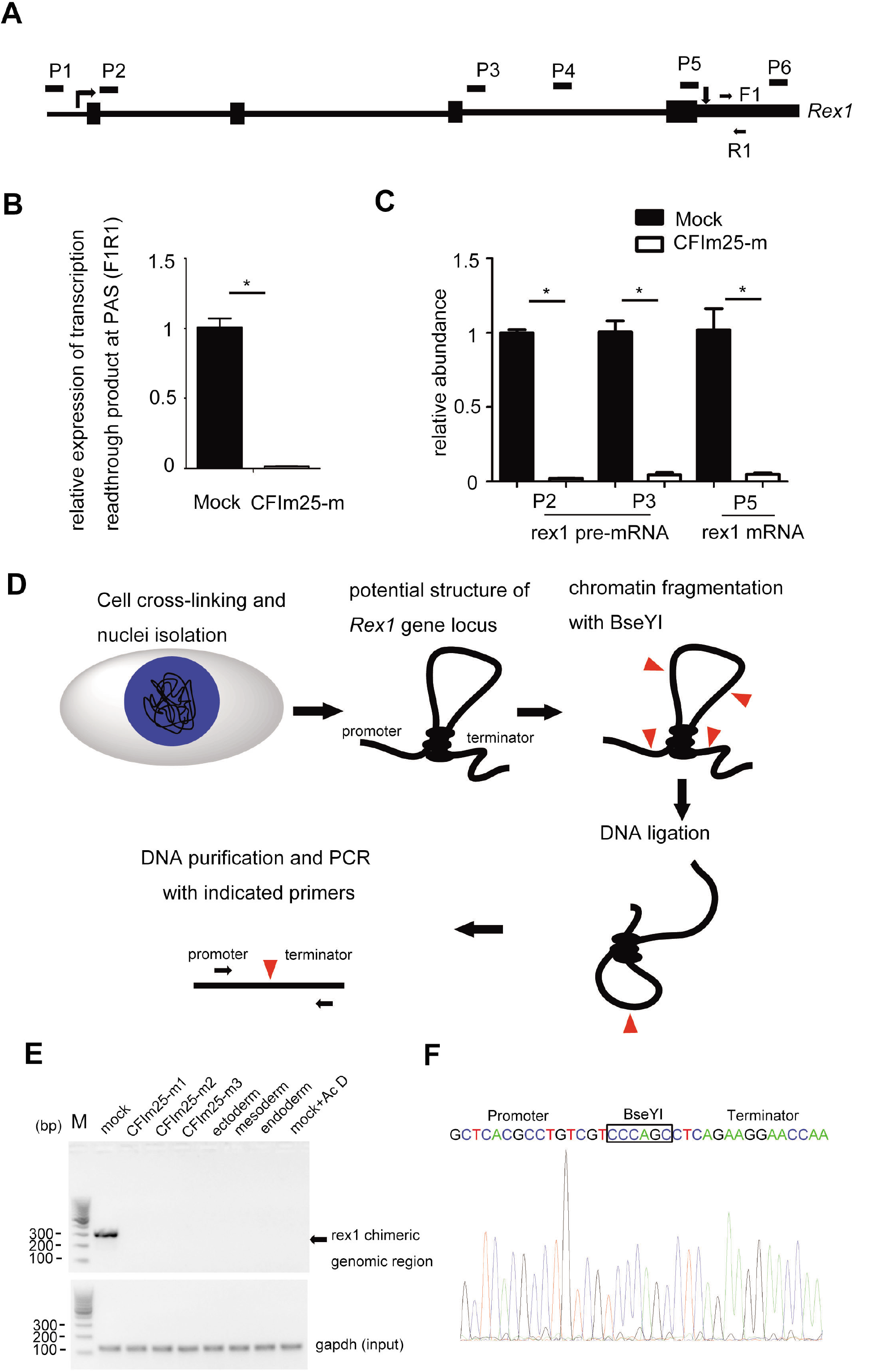
CFIm25 knockdown/mutation impacts rex1 gene transcription in H9 cells. (A-B) A pair of primers (F1/R1) was designed to detect the gene transcription readthrough beyond rex1 PAS. The relative expression of extended transcript in mock and CFIm25-m H9 cells was estimated by RT-qPCR analysis shown in Figure 3B. Gapdh gene expression serves as internal control. Student’s t-test was used to estimate the significance of the change. *p<0.05. (C) RT-qPCR analysis of the expression level of rex1 pre-mRNA and mRNA using indicated primers in mock and CFIm25-m cells. Student’s t-test was used to estimate the significance of the change. *P<0.05. (D) Outline of the 3C procedure used to detect chromatin interactions between promoter and terminator region for rex1 gene. (E) PCR product resulting from 3C library amplification using the primers located in the promoter and terminator regions. PCR product targeting gapdh gene serves as input. Cells used for 3C library preparation are indicated above the gel image. (F) Sanger sequencing shows that the PCR products correspond to the ligated DNA fragments of the two regions located at the promoter and terminator of rex1.

Given the essential role played by a promoter in gene transcription, we hypothesized that CFIm25 might be regulating activity of *rex1*’s promoter. To test this, we cloned *rex1* gene promoter into pGL3-basic plasmid, then measured its luciferase activity. Results showed no significant differences in luciferase activities between control and CFIm25-mutant cells (Figure 3-figure supplement 1D), suggesting that other elements might be involved in CFIm25’s role in regulating *rex1* transcription initiation/elongation.

Previous studies have shown that in yeast, some mRNA 3’ end processing factors may regulate transcription by bridging the interaction between the promoter and terminator regions of specific genes, a phenomenon termed gene looping (Al Husini *et al*., 2013; Allepuz-Fuster *et al*., 2019; Ansari & Hampsey, 2005; El Kaderi *et al*., 2009). In the present study, we used chromatin conformation capture (3C) analysis to test this model in *rex1* gene, based on following observations. Firstly, mouse *rex1* gene locus is characterized by long-range DNA-DNA interactions (Zhang *et al*, 2019). Secondly, ChIP-qPCR results suggested that CFIm25 was moderately enriched at both ends of the *rex1* gene in H9 cells (Figure 3-figure supplement 1E). Therefore, we constructed 3C libraries by digesting nuclei prepared from control and CFIm25-mutant cells with BseYI restriction enzyme. Then, we applied DNA ligation with T4 DNA ligase and PCR amplification targeting the indicated genomic sites (Figure 3D). BseYI enzyme was chosen as both *rex1* gene promoter and terminator region harbor this restriction enzyme site. Results revealed clear band, indicative of a genomic interaction between the *rex1* promoter and terminator regions, in control H9 cell lines, but not in CFIm25-mutant cells (Figure 3E; Figure 3-source file 1). However, the amount of DNA input was comparable in the parallel experiments. Result of sanger sequence analysis confirmed that the amplified PCR product were similar to those obtained near *rex1* promoter and terminator regions (Figure 3F).

Next, we evaluated whether the detected *rex1* promoter/terminator interaction was correlated with its expression. Strikingly, results from 3C-PCR analysis revealed no significant interaction in promoter/terminator interaction across differentiated cells expressing low levels of *rex1* transcript (Figure 3E; Figure 3-figure supplement 1F; Figure 3-source file 1), suggesting that this gene looping may be associated with gene expression. Similarly, treatment of the cells with transcription inhibitor Act D significantly abolished the observed interaction (Figure 3E; Figure 3-source file 1). Overall, these results indicated that CFIm25 might promote *rex1* expression in hESCs, in part, by facilitating formation of gene looping, a chromatin conformation status associated with transcription activation or enhancement.

### CFIm25 significantly affects gene transcription dynamics in hESCs

We hypothesized that CFIm25 could be promoting expression of other mRNA targets via transcriptional mechanisms. To this end, we performed RNA polymerase II (RNAPII) ChIP-seq in two mock control and CFIm25-mutant hESCs (the mutant for all the ChIP-seqs refers to m1 and m3, two biological replicates, unless otherwise noted), and found that CFIm25 globally promoted RNAPII’s association with actively transcribed genomic region, as evidenced by the metagene plots for all expressed genes (FPKM>1, 14274 genes based on mRNA-seq) and the group of down-regulated genes (Figure 4A; Figure 4-figure supplement 1A). This trend was pronounced for both high- and low-abundance genes (Figure 4-figure supplement 1A), suggesting that CFIm25-regulated gene transcription might be a general phenomenon. A common change, as shown in representative genome browser view of *gapdh* gene, involved a mild decrease in the signal at transcription start site (TSS) and gene body following gene editing of CFIm25 (Figure 4-figure supplement 1B). We noted that RNAPII ChIP-seq signal near TES decreased, rather than increased, in CFIm25-mutant cells (Figure 4A), further supporting the aforementioned conclusion that overall PAS processing efficiency was not affected, as inefficient PAS processing often leads to retarded transcription termination and RNAPII accumulation downstream of transcription end site (TES) (Nojima *et al*, 2015). Next, we performed peak calling by MACS2 and identified differential binding events using DiffBind package. As expected, we identified thousands of differential binding sites, with a majority of them located in intron (Figure 4B; Supplemental Table 5). Consistently, 22% of the genes showing expression level changes upon CFIm25 gene editing harbor differential RNAPII binding sites (Figure 4B). To validate the ChIP-seq results, we randomly selected nine regions, subjected them to ChIP-qPCR (three biological replicates), and observed consistent trend for most of the sites (Figure 4C). Thus, we presumed that CFIm25 might significantly regulate the expression of mRNAs at the transcription level. Additionally, the RNAPII binding on the group of up-regulated genes (310 genes) appeared to be less affected by CFIm25 gene editing (Figure 4-figure supplement 1A). However, this might not be significant, because up-regulated genes intrinsically require more RNAPII binding to produce more transcripts.

**Figure 4.**
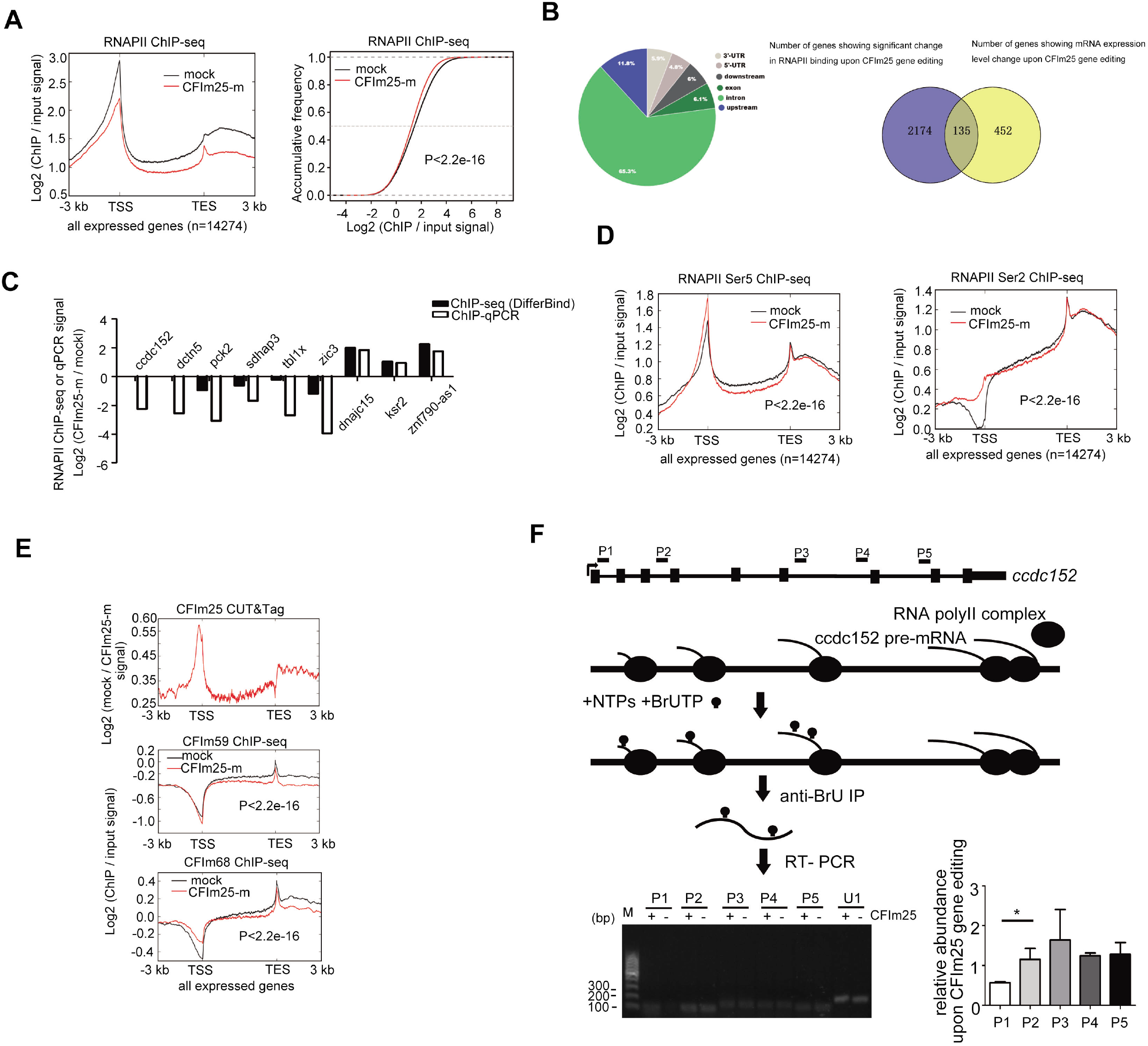
CFIm25 knockdown/mutation globally impacts transcription dynamics in H9 cells. (A) (left panel) Metagene plots of RNAPII ChIP-seq reads in mock and CFIm25-m H9 cells for actively transcribed genes (FPKM>1 based on mRNA-seq, 14274 genes in total), and its corresponding cumulative frequency plot (right panel). K-S test was used to examine the significance of the difference between the two plots. (B) (left panel) Pie plot showing the genomic annotations of 4024 sites that displayed differential RNAPII binding. Peak calling was performed using MACS2 software and DiffBind package was used to identify the differential binding events. (right panel) Venn diagram showing the numbers of overlapping and non-overlapping genes that displayed differential RNAPII binding and mRNA expression level change. (C) Comparison of RNAPII ChIP-seq and ChIP-qPCR results in mock and CFIm25-m H9 cells for the tested genomic sites. Y axis represents the average fold changes from replicates (ChIP-seq: two replicates; ChIP-qPCR: three replicates). (D) Metagene plots of RNAPII Ser5 ChIP-seq and RNAPII Ser2 ChIP-seq reads for actively transcribed genes in mock and CFIm25-m H9 cells. K-S test was used to examine the significance of the difference between the two plots. (E) Metagene plots of CFIm25 CUT-Tag (top), CFIm59 ChIP-seq (middle), and CFIm68 ChIP-seq (bottom) reads for actively expressed genes in mock and CFIm25-m H9 cells. CFIm25 CUT-Taq profile in CFIm25-m cells was used as normalization control. For CFIm59 and CFIm68 ChIP-seq, K-S test was used to examine the significance of the difference between the two plots. (F) Nuclear run-on assay on the nascent ccdc152 transcript. The gene structure and the primer positions are indicated on the top. The diagram for the nuclear run-on assay is shown in the middle. A representative set of RT-PCR data are shown in the bottom panel. Left gel image: nuclear run on assays followed RT-PCR using primers targeting P1-P5 region. ‘CFIm25-’ represents CFIm25-m cell nuclei, whereas ‘CFIm25 +’ represents mock cell nuclei. Right Bar graph represents RT-qPCR data from three independent experiments. U1 snRNA was assayed as normalization control. Student’s t-test was used to estimate the significance of the change. *P<0.05.

Given that *rex1* is a transcription-related pluripotency factor, whose expression is down-regulated in CFIm25-mutant cells, we further investigated whether the aberrant RNAPII occupancy might be caused by its depletion. Here, we generated a stable cell line expressing *rex1* shRNA, and subsequently analyzed it using RNAPII ChIP-seq assay. Results from qRT-PCR and western blot analysis revealed that *rex1* was moderately depleted (Figure 4-figure supplement 1C; Figure 4-source file 1). Significantly, rather than detecting a decrease, we observed an increase in RNAPII ChIP-seq signal in *rex1* RNAi cells (Figure 4-figure supplement 1C). Furthermore, the same bioinformatics pipeline and statistical analysis revealed no presence of differential binding sites, which is in contrast with results from RNAPII ChIP-seq in CFIm25-mutant cells (Supplemental Table 5). Additionally, we observed that other CFIm25 mRNA targets (84 down-regulated genes and 129 up-regulated genes upon CFIm25 gene editing) exhibited no clear general transcription-associated molecular functions (Supplemental Table 3). Thus, we combined these observations with the aforementioned data in which CFIm25 overexpression rescued the gene expression phenotype, and concluded that CFIm25 may be directly responsible for the observed transcription effect.

To further validate this finding, we performed ChIP-seq analysis using antibodies against RNAPII Ser5, a modification status associated with transcription initiation/elongation (Hsin & Manley, 2012; Lyons *et al*, 2020). Consistent with the finding from previous studies that RNAPII Ser5 has major binding peak at TSS, our results revealed similar patterns in hESCs. Results from both metagene plot and differential peak identification revealed potential transcription initiation/elongation disturbance at the transcriptome level (Figure 4D; Figure 4-figure supplement 1D; Supplemental Table 5). Additionally, we performed RNAPII Ser2 ChIP-seq analysis. Previous studies have shown that RNAPII Ser2 interacts with transcription termination and its signal gradually increases toward TES (Tellier *et al*, 2020). Here, we observed similar binding patterns in hESCs, confirming reliability of our data (Figure 4D). Moreover, the level of Ser2 signal showed little, if any, change at TES region, thereby supporting the conclusion that CFIm25 mutation does not markedly affect the overall efficiency of transcription termination. The genome browser views for two representative genes (down-regulated upon CFIm25 gene editing) were shown in Figure 4-figure supplement 1E respectively.

To identify additional evidence supporting CFIm25’s role in regulating gene transcription, we performed CUT&Tag analysis using antibodies against CFIm25. Strikingly, in addition to the peak observed at TES, we detected a sharp peak at TSS in the CFIm25, but not CFIm68 or CFIm59, binding profiles (Figure 4E). The positive CFIm25 binding signal throughout the gene body for all expressed genes, including the group of down-regulated/up-regulated genes (Figure 4-figure supplement 1F), provides another line of evidence that CFIm25 might play a general role in gene transcription regulation relative to its CFIm counterparts. Notably, although results from both CUT&Tag and ChIP-seq analyses are often limited by non-specificity of target antibodies, our findings in CFIm25 are reliable owing to prior normalization of the signals by backgrounds from CFIm25-mutant cells.

Furthermore, we performed a nuclear run-on assay, which provides a measure of transcription and minimizes the effect of RNA stability, and analyzed expression levels of two target pre-mRNAs by qRT-PCR, these two genes were selected owing to their differential mRNA expression as well as effect on their transcription processes as shown by RNAPII/Ser5 ChIP-seq and RNAPII ChIP-qPCR results (Figure 4C; Figure 4-figure supplement 1E). Notably, we detected low levels of transcription product near the promoter region in CFIm25-mutant cells, but not in the middle or at the end point of these genes (Figure 4F; Figure 4-figure supplement 1G; Figure 4-source file 2). Taken together, these results suggest that CFIm25 may be playing a cellular role in the early stages of gene transcription in specific genes, including *ccdc152* and *dctn5*.

To validate our findings, we analyzed a randomly selected dataset from a previous gene expression dataset in which CFIm25 was depleted in human cancer cell line (Routh *et al*, 2017). In addition to APA change, we observed differential expression of thousands of transcripts upon CFIm25 depletion (Figure 4-figure supplement 1H; Supplemental Table 6). Interestingly, these two groups of genes did not show a striking overlap, further indicating that CFIm25 might have functions other than PAS usage.

### CFIm25 might regulate transcription through its association with LEO1

Next, we explored the mechanism through which CFIm25 regulates transcription. Since some splicing factors could regulate transcription process through their interaction with general transcription factors (Caizzi *et al*, 2021; Lin *et al*, 2008), we hypothesized that CFIm25 might also utilize such a mechanism to regulate transcription. To this end, we took advantage of the aformentioned 3XFlag-CFIm25 H9 cell line and performed anti-FLAG immunoprecipitation (IP) assays followed by mass spectrometry (MS) analysis. As expected, CFIm68 and CFIm59, two known CFIm25 interaction partners, were highly enriched in the FLAG IP sample based on cell lysates prepared from hESCs overexpressing FLAG-CFIm25 (Supplemental Table 7). LEO1, an RNAPII associated factor, was selected among the candidates owing to its direct association with transcription (Xie *et al*, 2018).

MS results were further confirmed by western blot analysis (Figure 5A; Figure 5-source file 1). As our FLAG IP/MS was carried out in the absence of ribonuclease, it is possible that the detected interaction was mediated by RNAs. To confirm a direct protein-protein interaction, we performed aformentioned FLAG IP in the presence of RNAse A to avoid RNA-mediated effects. Western blot analysis revealed similar result to that without RNAse A treatment, indicating a direct association of CFIm25 with LEO1 (Figure 5-figure supplement 1A; Figure 5-source file 1). To further confirm this, we carried out GST-pull down assays using recombinant GST-tagged CFIm25 protein and His-tagged LEO1 protein (Figure 5-figure supplement 1B; Figure 5-source file 2). A series of LEO1 truncation proteins were used as we failed to obtain the full length LEO1. Significantly, we observed that LEO1 C-terminus fragment, but not other fragments (truncation fragment 4 and 5 were apparently detectable in western blotting analysis using the anti-His Tag antibody, whereas their expression were not readily detectable using Coomassie blue staining), showed detectable association with CFIm25 under physiological conditions, as shown by the western blot analysis (Figure 5B; Figure 5-source file 2).

**Figure 5.**
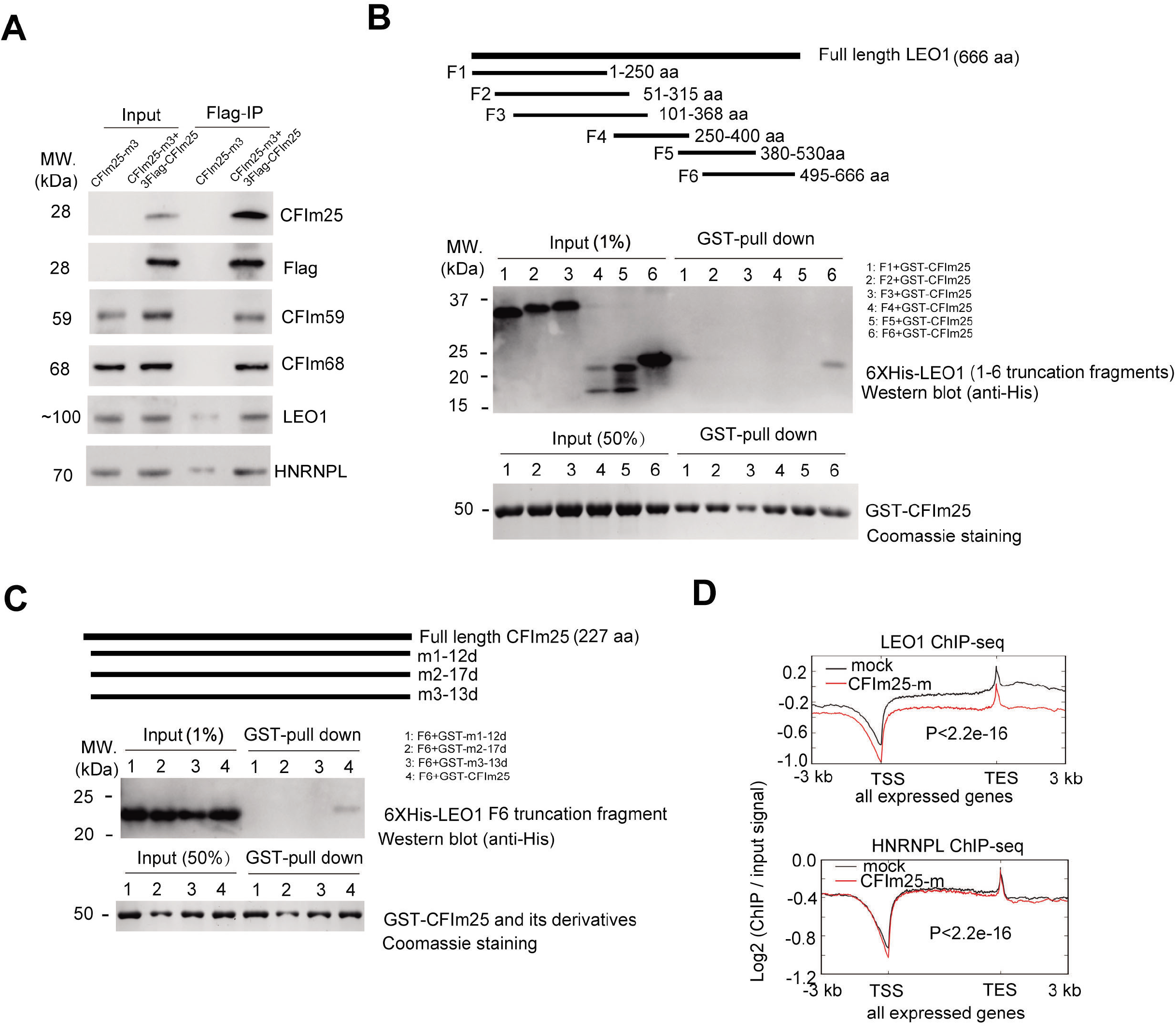
CFIm25 interacts with LEO1 and impacts the DNA genomic binding profile of LEO1. (A) Western blot analysis of the abundance of indicated proteins in Flag-IPed sample. Flag-IP was performed using extracts from CFIm25-m3 and CFIm25 (m3+3XFIag-CFIm25 overexpression) H9 cells. Input: 1% of the lysates for IP. The primary antibody for CFIm25 is from santa cruz (sc-81109). (B) Schematic representation of full length human LEO1 protein and truncation fragements (F1-F6) (upper). Truncation fragments were fused to pET-28a vector (BamHI and XhoI) for recombinant His-tag protein expression. At least three independent experiments have been performed, and a representative western blotting result of GST-pull down assay using anti-His antibody is shown in the middle picture. As the bait protein, recombinant GST-CFIm25 was stained with Colloidal Coomassie G-250 (Bottom). The percentage of input is indicated in the bracket. (C) Schematic representation of human CFIm25 protein and its N-terminus deletion/mutation derivatives m1-12d/m2-17d/m3-13d represent the 12/17/13 amino acids deletion/mutation proteins produced in CFIm25-mutant (m1-m3) ceils respectively) (upper). Full length and mutant CFIm25 proteins were fused to pGEX-4T3 vector (BamHI and XhoI) for recombinant GST-tag protein expression. In the GST pull-down assay, recombinant His-tag LEO1-F6 protein was used as the prey protein. At least three independent experiments have been performed, and a representative western blotting result of GST-pull down assay using anti-His antibody is shown in the middle picture. GST-fused bait proteins are stained with Colloidal Coomassie G-250 (Bottom). The percentage of input is indicated in the bracket. (D) Metagene plots of LEO1 ChIP-seq and HNRNPL ChIP-seq reads for actively expressed genes in mock and CFIm25-m cells. K-S test was used to examine the significance of the difference between the two plots.

Inspired by the above result, we further tested the associations of LEO1 C-terminus fragment with several CFIm25 N-terminus mutants. Three mutants were designed to mimic the three small N-terminus deletion/mutant proteins produced by CFIm25 gene-edited cells (Figure 1-figure supplement 1B; Figure 5-figure supplement 1B; Figure 5C). Strikingly, we observed that none of these 3 mutants were able to associate with LEO1 C-terminus truncation fragment, in comparison with wild type GST-CFIm25 (Figure 5C), providing evidence that CFIm25 may associate with LEO1 through its N-terminus, and the transcription phenotype in CFIm25-mutant cells might be caused by the absence of this protein-protein interaction. It must be noted, nevertheless, that this interaction is relatively weak in vitro based on the pull-down efficiency (Figure 5B, 5C).

To explore the functional impact of CFIm25-LEO1 association, we carried out ChIP-seq analysis on LEO1 in CFIm25-mutant alongside control hESCs, and found that LEO1 exhibited a significant decrease in binding frequency on transcribed genes, including the group of down-regulated genes, upon CFIm25 gene editing (Figure 5D), although the overall ChIP efficiency is relatively lower than that of RNAPII (Figure 4A and 5D). Interestingly, the decrease trend seemed more obvious for high-abundance genes (Figure 5-figure supplement 1C), as shown in representative genome browser view of *gapdh* gene (Figure 5-figure supplement 1D). This observation is in agreement with aforementioned finding that RNAPII occupancy is globally down-regulated in CFIm25-mutant hESCs (Figure 4A). It is important to point out that the input samples gave approximately the same signal in our RNAPII and LEO1 ChIP-seqs (Figure 4-figure supplement 1B; Figure 5-figure supplement 1D), and thus the detected discrepancy between ChIP samples did not appear to be caused by DNA heterogeneity in input samples. In contrast, the overall DNA binding pattern of HNRNPL, another protein that has potential interaction with CFIm25 (Figure 5A; Supplemental Table 7; Figure 5-source file 1), showed little, if any, change in CFIm25-mutant cells (Figure 5D). Taken together, these results suggest that CFIm25 potentially affects the genomic binding pattern of its associated transcription factor LEO1, thereby providing a potential mechanism underlying CFIm25-mediated transcription regulation.

### CFIm25 targets associate with the phenotypes of CFIm25 gene editing in hESCs

The above results suggest that CFIm25 may affect gene transcription process and enhance expression of a subset of mRNA targets. To understand the effect of CFIm25 gene editing on cellular phenotypes, we used overexpression and knockdown experiments on several high-confidence CFIm25 targets, then analyzed the resulting cellular phenotypes. Strikingly, depletion of *rex1* significantly impaired differentiation of the endoderm in hESCs, as evidenced by downregulation of endoderm-lineage markers following differentiation induction (Figure 6A). Notably, we found no significant changes in expression of most of the tested self-renewal markers and the cell morphology in both CFIm25-mutant and *rex1* RNAi cells (Figure 6-figure supplement 1A and B), suggesting that *rex1* depletion did not affect self-renewal of H9 cells. These results are consistent with the findings of previous reports in which mouse *rex1* was reportedly dispensable for self-renewal of ES cells (Masui *et al*., 2008). In fact, knocking it out in mouse ES cells was implicated in impaired differentiation of the visceral endoderm (Masui *et al*., 2008). Furthermore, we carried out *rex1* gene overexpression in *rex1* RNAi hESCs and performed parallel experiments. Indeed, *rex1* overexpression could partially rescue the differentiation potential phenotype induced by *rex1* depletion (Figure 6A).

**Figure 6.**
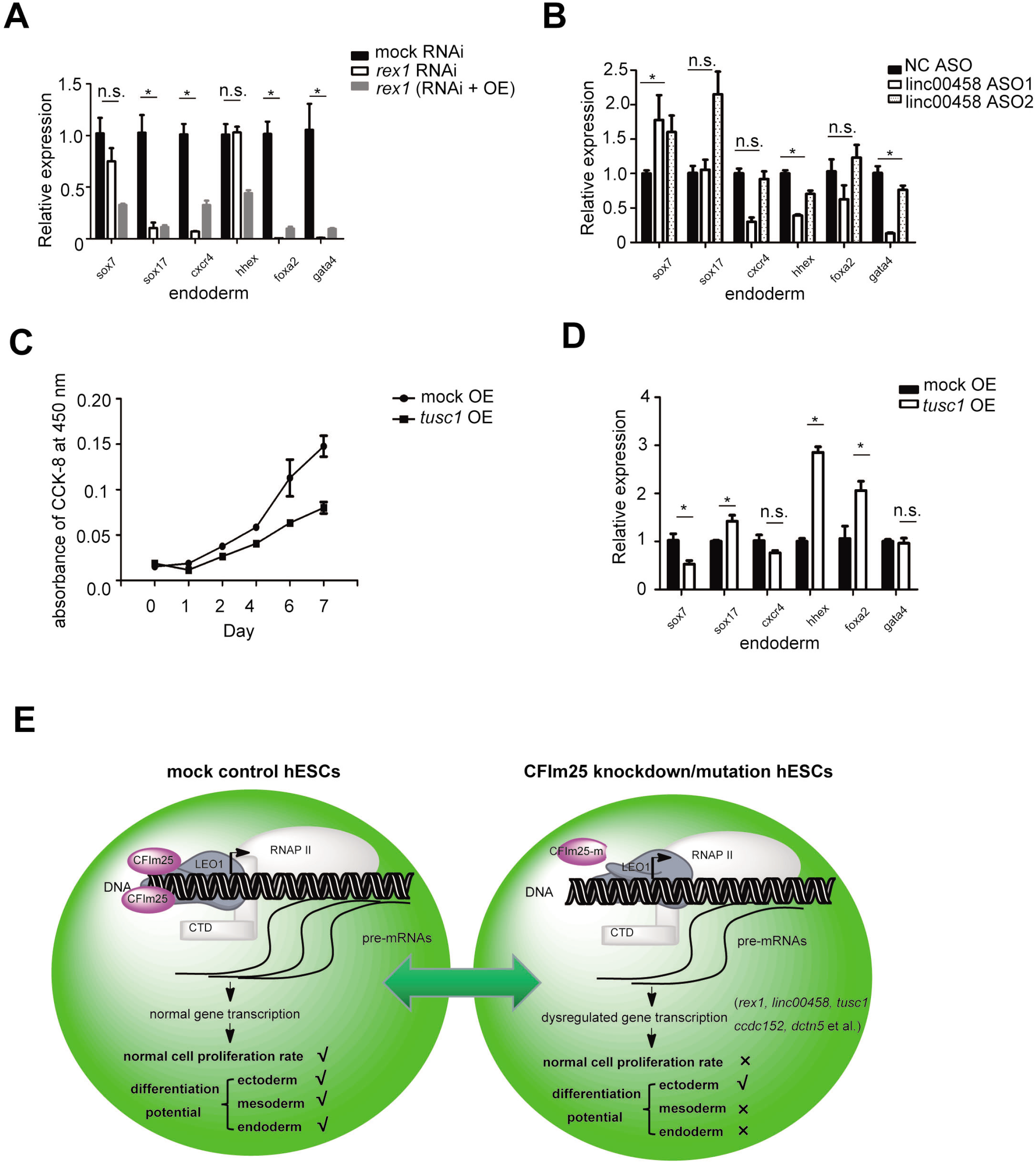
CFIm25 targets play roles in hESCs cell proliferation and pluripotency. (A-B; D) RT-qPCR analysis of the expression level of 6 endoderm lineage differentiation markers in indicated cell lines during endoderm lineage differentiation (OE: overexpression; ASO: antisense oligo). Three independent experiments have been performed and quantified results are shown. Student’s t-test was used to estimate the significance of the change. *P<0.05; ns: non-significant. (C) Cell proliferation rate measurement by CCK-8 kit for mock and tusc1 gene overexpression H9 cell lines. Three independent experiments have been performed and quantified results are shown. (E) A schematic model summarizing the key finding in this study. Mutations in human CFIm25 protein N-terminus did not significantly affect cellular mRNA alternative polyadenylation profile, but rather affected transcription process, thereby decreasing the expression level of a group of transcripts associated with pluripotency (such as rex1 gene) and cell proliferation (such as tusc1 gene). CFIm25 depletion/mutation predominantly caused defects in the endoderm/mesoderm differentiation and accelerated the rate of cell growth in H9 cells.

Since *rex1* depletion could not fully recapitulate the endoderm lineage differentiation phenotype caused by CFIm25 gene editing (Figure 1D and 6A), we tested the function of another CFIm25 high-confidence target, *linc00458*, a long noncoding RNA that has been associated with endodermal lineage specification (Chen *et al*, 2020). Results showed that *linc00458* knockdown using Antisense Oligonucleotides (ASO) technology significantly down-regulated endoderm-specific genes *gata4* and *hhex* during induction of endoderm differentiation (Figure 6B). Overall, these results suggest that the observed phenotype in hESCs lacking CFIm25 might be caused by synergistic effects of CFIm25 mutation in target genes.

We further tested the function of several other targets that might be associated with cell proliferation phenotype upon CFIm25 gene editing. As expected, overexpression of *tusc1*, a tumor-associated suppressor gene (Shan *et al*, 2013), caused an apparent suppression in the rate of cell proliferation (Figure 6C). When cells overexpressing tusc1 gene were subjected to endoderm differentiation, they appeared to be dysregulated during this process, as evidenced by the expression of molecular markers (Figure 6D). This result was consistent with the findings from previous reports in which some tumor suppressor genes were found to play a crucial role in ESCs pluripotency (Fu *et al*, 2020; Langer *et al*, 2019). Taken together, our results indicate that phenotypes of CFIm25-mutant hESCs result from down-regulation of a subset of CFIm25-regulated RNA transcripts.

## Discussion

In the field of co-transcriptional mRNA processing, most previous reports studying co-transcriptional mRNA 3’ processing have focused on how transcription facilitates mRNA 3’ processing, and the effect of 3’ processing on mRNA alternative polyadenylation (APA). In this study, we present evidence that CFIm25, a canonical mRNA 3’ processing factor, may promote gene transcription in H9 cell line, and the mechanism might be involved in its interaction with LEO1, an RNAPII associated factor. Importantly, CFIm25 as well as its targets plays a direct role in H9 cell function. A schematic model is presented in Figure 6E. Our findings not only provide novel insights into the critical role played by CFIm25 (and possibly other 3’ processing factors) in gene regulation, aside from its traditionally studied function in mRNA 3’ processing and APA regulation, but also expand our understanding of its role in determination of cell fate.

Researchers have long hypothesized that mRNA 3’ processing factors may be playing a role in transcription. For example, the co-purification of CPSF with TFIID was discovered more than twenty years ago (Dantonel *et al*., 1997). Recent studies have shown that CstF64 and CPSF73 regulate RNAPII activity at transcription end sites (TES) (Nojima *et al*., 2015), and CFIm25/CFIm68 depletion in HeLa cells affects RNAPII occupancy in a subset of genes (Tellier *et al*., 2018, 2019). However, our results are significant in at least two major respects. Firstly, we excluded the possibility that the observed transcription phenotypes might be caused by impaired transcription termination, upon CFIm25 gene editing. Therefore, our findings provide more direct evidence that mRNA 3’ processing factor may be playing an active role in early transcription rather than passively interacting with transcription termination. Secondly, results from our global analyses and nuclear run-on assays for specific genes affirm reliability of our results, while the findings of our CFIm25 overexpression rescue experiment validate the conclusion.

Given the global effect of RNAPII occupancy on transcribed genes in CFIm25-mutant cells, it remains unclear why CFIm25 gene editing only affected the steady level of a specific subset of genes, as we observed no significant difference in total poly(A+) RNA yield between control and CFIm25-mutant hESCs (Figure 6-figure supplement 1C). We attribute this to two scenarios. Firstly, the steady levels of mRNAs are controlled by multiple factors, such as transcription, mRNA processing and stability (Slobodin *et al*, 2020), while we cannot rule out existence of unknown mechanisms that regulate this balance in mRNA expression upon CFIm25 gene editing. For example, we noted a slight increase, albeit statistically insignificant, in canonical SVL PAS processing efficiency (Figure 2-figure supplement 1B). Therefore, it is plausible that the steady levels in a majority of genes with no apparent change in expression might be balanced by decreased transcription and increase in 3’ processing efficiency. Secondly, transcription itself is controlled by auto-regulatory mechanisms. For example, paused RNAPII reportedly inhibits new transcriptional initiation (Shao & Zeitlinger, 2017). In the present study, we used RNAPII ChIP-seq analysis to reveal defects in the observed global transcription. However, the extent to which the occupancy of RNAPII contribute to the transcription output in our system remain unknown.

Further studies are required to fully understand the role of CFIm25 in transcriptional regulation in the context of co-transcriptional mRNA processing. Firstly, although it is unlikely that this phenomenon is unique to hESCs, we cannot fully exclude this possibility. Similar assays in other cell types are imperative to validate these findings and unravel the precise underlying molecular mechanisms. Secondly, previous studies have shown that CFIm25 can regulate global mRNA alternative polyadenylation (APA) in many cell types (Alcott *et al*., 2020; Brumbaugh *et al*., 2018; Chu *et al*., 2019; Huang *et al*., 2018; Jafari Najaf Abadi *et al*., 2019; Masamha *et al*., 2014; Tan *et al*., 2018; Weng *et al*., 2019; Zhu *et al*., 2018), while recent reports demonstrated that it could also regulate mRNA splicing in specific genes (Gao *et al*, 2020; Scarborough *et al*, 2021). Future explorations are expected to reveal whether they are associated with CFIm25’s potential role in transcriptional regulation, and to elucidate mechanisms underlying coordination of these multiple regulatory roles. Finally, we envisage that further explorations will generate a deeper understanding of the functional significance of CFIm25-mediated regulation of transcription. Previous studies have shown that CFIm25 plays important cellular roles under normal physiological conditions, while its dysregulation has been associated with a variety of diseases, such as cancer, learning deficits and dermal fibrosis (Alcott *et al*., 2020; Brumbaugh *et al*., 2018; Chu *et al*., 2019; Huang *et al*., 2018; Jafari Najaf Abadi *et al*., 2019; Masamha *et al*., 2014; Tan *et al*., 2018; Weng *et al*., 2019). Results of the present study corroborated the aforementioned findings, as evidenced by enhanced cell proliferation and impaired differentiation potential in hESCs upon CFIm25 mutation. It is plausible that CFIm25-mediated transcription regulation may also be involved in other reported cellular systems. A key challenge for future investigations is emergence of multiple molecular functions of CFIm25. For example, although CFIm25 might regulate mRNA abundance, splicing and APA for the same group of genes, approaches for delineating their respective contributions to cellular phenotype remain limited. With the growing trend in generating related data, we believe a clearer picture will be painted with regards to the functional significance of CFIm25-mediated regulation in transcription.

## Materials and Methods

### Cell culture and plasmids transfections

H9 hESCs were purchased from the National Collection of Authenticated Cell Cultures (Shanghai, Catalog SCSP-302) and were maintained in mTeSR (Stem Cell Technology) on Matrigel-coated plates at 37°C. CRISPR/Cas9-mediated CFIm25 gene editing was carried out using a previously reported eCRISPR plasmid (Xie *et al*., 2017), based on the following gRNA target sequences: g1: CAGCCGGTCTGCGAGCGATT, g2: CCGAACTGAGTGACCCCCCG, and g3: CCAATCGCTCGCAGACCGGC. Cultures were selected on puromycin, single-cell clones picked for further expansion. pLKO.puro shRNA vectors were used for *rex1* RNAi, with target sequence GCATGCAAATACGAACAAGAA, while lentiviral plasmids were used for gene overexpression. Briefly, 3XFlag-CFIm25 cDNA was cloned into CD533A-2 pCDH-EF1-MCS-IRES-Neo (SBS), whereas *rex1* and *tusc1* overexpression plasmids were purchased from Fulengen, Catalogs EX-T4815-Lv242; EX-I2275-Lv233. Transfections for plasmids and Antisense oligos (ASOs) were performed using Lipofectamine 3000 and Lipofectamine RNAiMAX (Life Technology), respectively. ASOs targeting *linc00458* were ordered from RiboBio. Cells were harvested at the suggested time points for further analysis upon transfection.

### Cell growth measurement and differentiation induction

Cell growth monitoring and analysis of hESCs trilineage differentiation were performed using the Cell Counting Kit-8 (Dojindo) and STEMdiffTM trilineage differentiation kits (Stem Cell Technology), according to the manufacture’s protocols. For cell growth measurement, cells were seeded at 5000 cells per well on 96-well plate at day 0. After the addition of CCK-8 solution, the absorbance at 450 nm was measured using a microplate reader at the indicated time points (day 0, 1, 2, et al.). It is important to note that the growth rate of human embryonic stem cells is sensitive to the quality and density of starting cells. Therefore, it is essential to keep the starting cell numbers at the same level and make sure tested cells were treated in parallel in this experiment. For stem cell trilineage differentiation induction experiments, cells were seeded in 12-well plate at the suggested cell density. The induction time is approximately within one week. After induction, total RNAs were harvested, and subsequent RT-qPCR analysis of lineage expression markers was carried out to estimate the induction efficiency. Moreover, differentiation of hESCs cardiomyocytes was performed using the STEMdiffTM Cardiomyocyte Differentiation Kit (Stem Cell Technology), whereas analysis of cardiomyocyte induction efficiency was conducted via FACS using the cTnT+ primary antibody (Thermo Scientific, MA5-12960).

### Luciferase reporter assays

hESCs were transfected for 24 h with pPASPORT-SVL PAS or pGL3-basic (promoter sequence inserts)+pRL-TK plasmids, harvested, then subjected to analysis of Luciferase activity using the Promega Dual-Luciferase Reporter kit and Beirthold Sirius detection system.

### RNA-biotin based pull-down assay

SVL PAS RNA and the corresponding point mutant RNA (CPSF recognition motif ‘AAUAAA’ hexamer was mutated to ‘AACAAA’) were made by in vitro transcription using SP6 polymerase, and biotinylated at 3’ end using a biotinylation Kit (Thermofisher). H9 cell nuclear extracts (NEs) were made following the described protocol (Huang *et al*, 2017; Shi *et al*., 2009). Approximately 15 μg biotinylated RNAs were first bound to the streptavidin beads, and then incubated with 100 μl pre-cleared NE in the polyadenylation condition [40% NE, 8.8 mM HEPES (pH 7.9), 44 mM KCl, 0.4 mM DTT, 0.7 mM MgCl2, 1 mM ATP, and 20 mM creatine phosphate] for 20 minutes, after biotin-streptavidine binding, washing, pull-down sample were heated (75°C for 5 minutes) in 1XSSC buffer (150 mM NaCl, 15 mM sodium citrate) for elution. The eluted sample was further subjected to western blot analysis.

### Metabolic Pulse-Chase RNA Labeling with bromouridine (BrU)

Cellular pre-mRNA labeling was performed with bromouridine, according to a published protocol (Paulsen *et al*, 2014). Briefly, hESCs were grown to approximately 50% confluency in 3 10-cm plates, then incubated with bromouridine (final 2 mM), at a pulse time of 30 min. BrU containing pre-mRNA was purified with 2 μg anti-BrdU antibodies (BD Pharmingen) prior to use in downstream RT-qPCR analysis.

### Nuclear run-on assay

Nuclear run-on assays were performed using previously described protocol with minor modifications (Lin *et al*., 2008; Roberts *et al*, 2015). Briefly, 1 × 10^7^ hESCs were permeabilized with digitonin, and nuclei was isolated via low-speed centrifugation. A nuclear run-on reaction was initiated by mixing the nuclei with 60 μl reaction buffer (50 mM Tris-HCl, pH7.4, 10 mM MgCl2, 150 mM NaCl, 25% (v/v) glycerol, 0.5 mM PMSF and 25 U ml-1 RNasin) and 40 μl BrU-containing NTPs mixture (1.8 mM ATP, 0.5 mM CTP and GTP, 0.375 mM UTP, 0.125 mM BrU), with a 15-min incubation at 25°C. After the reaction, RNA was extracted from BrU-containing cells using the Trizol reagent (Thermo Scientific), and further isolated by 2 μg anti-BrU antibodies (BD Pharmingen). Purified RNAs were used for downstream RT-qPCR analysis.

### Chromosome Conformation Capture (3C) analysis

A 3C analysis was carried out according to a published protocol (El Kaderi *et al*, 2012), with minor modifications. Briefly, 1 × 10^7^ cells were cross-linked in 1% formaldehyde solution and quenched with 125 mM glycine. Cells were permeabilized, their nuclei were isolated via centrifugation then resuspended in a 0.5 ml solution comprising 1.2 × restriction enzyme NEBuffer™ r3.1 (NEB) containing 0.3% SDS. After incubation, shaking, and digestion with BseYI enzyme (NEB), the cross-linked chromatin was ligated using T4 DNA ligase (NEB). The DNA was de-crosslinked, purified via phenol/chloroform extraction and ethanol precipitation, then subjected to 3C-PCR analysis using primers listed in supplementary table 8. PCR reactions were set up by mixing: 500 ng of DNA template; 25 pmol of each primer; 5 μl of 10× PCR buffer; 1 μl of 10 mM dNTP mix; 1 μl of 5 U/μl taq DNA polymerase; H2O to bring the final volume to 50 μl. Run PCR with the following thermal cycling parameters.1 cycle: 2 min 95°C (initial denaturation); 30 cycles: 30 sec 95°C (denaturation), 30 sec 55°C (annealing), 1 min 72°C (extension); 1 cycle: 4 min 72°C (final extension).

### 3’-seq, mRNA-seq, ChIP-seq

We performed 3’-seq analysis using QuantSeq Rev 3’ mRNA sequencing library prep kit (Lexogen), on the NovaSeq platform. Raw reads were reverse complemented and mapped to the human genome (hg19), allowing up to two mismatches using Bowtie2 with the settings ‘bowtie2 -p 28 -N 1 -k 1’. The 3’ end of the read maps was considered a poly(A) junction. The bioinformatics analysis for reads filtering and clustering, internal priming removal, poly(A) site identification and subsequent APA analysis shown in Figure 2A and Figure 4-figure supplement 1H, were performed essentially as previously described (Lackford *et al*., 2014; Yao *et al*., 2012).

Preparation of mRNA-seq library, sequencing and analysis of sequence data were performed in accordance with the standard protocol described by Illumina and Novogene. Identification of differentially expressed genes was done using the DESeq2 tool.

ChIP-seq libraries were prepared using the ChIP-IT® Express Enzymatic Shearing (Active Motif) and ChIP-seq library preparation (Vazyme) kits. Primary antibodies used for ChIP included RNAPII (39497, Active Motif); hnRNPL (18354-1-AP, Proteintech); RNAPII Ser5 (61986, Active Motif); RNAPII Ser2 (61984, Active Motif); LEO1 (PAB14102, Abnova); CFIm68 (A301-358A, Bethyl); CFIm59 (A301-359A, Bethyl). ChIP-seq libraries for CFIm25 were prepared using the CUT & Tag Hyperactive In-Situ ChIP Library Prep Kit (Vazyme) and primary antibody (10322-1-AP, Proteintech). These libraries were sequenced on the NovaSeq platform, and all ChIP-seq data processing and analysis performed according to the ENCODE ChIP-seq pipeline (https://www.encodeproject.org/data-standards/chip-seq/). Metagene plots were generated with Deeptools2 computeMatrix tool with a bin size of 50 bp and plotProfile –outFileNameData tool. Graphs representing the (IP/Input) signal (ChIP-seq) were then created with R packages. Metagene profiles are shown as the average of two biological replicates. P-values were computed with a Kolmogorov-Smirnov (K-S) test.

### Affinity purification of CFIm25-associated proteins

A total of 10×10^7^ hESCs cells that stably overexpress Flag-tag CFIm25 or negative control were harvested by centrifugation at 2000 g for 5 min. Cells were lysed with 3 ml IP lysis buffer (87787, Thermo Scientific) in the presence of protease inhibitor cocktail (Roche). After incubation at 4 °C for 20 min and centrifugation at 15000 g for 10 min, cell extracts (3 ml of the supernatant) were incubated with Anti-Flag Affinity Gel (Bimake) at 4 °C for 3 h. After three washes, each with 1 ml Wash Buffer (30 mM Tris-Cl, pH 7.4, 150 mM NaCl, and 0.5% Triton X-100), proteins were eluted from the beads using elution buffer (30 mM Tris-Cl, pH 7.4, 150 mM NaCl, 0.5% Triton X-100, and 400 μg/ml Poly FLAG peptide). Eluted samples were resolved in an SDS-polyacrylamide gel, followed by mass spectrometry (Mass Spectrometry Facility at Novogene, Beijing). Aliquots of the eluted proteins were used for western blotting.

### GST pull-down assay

Human CFIm25 and corresponding N-terminus mutants were inserted into vector pGEX-4T3 and expressed as GST-CFIm25 fusion protein in BL21 (DE3) strain. The fusion protein was purified with ProteinIso® GST Resin (TRANS). LEO1 truncation fragments were inserted into vector pET-28a vector and expressed as His-LEO1 fusion proteins in BL21 (DE3) strain. The fusion proteins were purified with HisPur Cobalt Resin (Thermo Scientific). For GST pull-down assay, two proteins (approximately 10 μM for each) were mixed in binding buffer (20 mM HEPES, pH 7.9, 150 mM NaCl, 1 mM MgCl2, 0.2 mM EDTA, 0.1% NP-40, protease inhibitor). After binding, washing, proteins were eluted from the beads using elution buffer (20 mM HEPES, pH 7.9, 150 mM NaCl, 1 mM MgCl2, 0.2 mM EDTA, 0.1% NP-40, protease inhibitor, 20 mM Glutathione). Eluted proteins were used for western blotting or Coomassie blue staining.

### Quantitative real time polymerase chain reaction (qRT-PCR) and western blot analysis

Quantitative real-time PCR was performed in 96-well plates, on the LightCycler® 480 qPCR system (Roche). Briefly, RNAs were quantified on a NanoDrop™ 1000 Spectrophotometer (Anti-BrU antibodies purified pre-mRNAs were not quantified due to low yield). The cDNA was synthesized from extracted RNA using the superscript III reverse transcriptase kit (Life Technology). The cDNA was used for qRT-PCR amplification targeting genes outlined in Supplementary Table 8. Expression data were analyzed using the ΔΔCt method, and normalized based on appropriate controls. All the qPCR parameters and results including reaction conditions, input volumes and Ct values, have been listed in MIQE form as Supplemental Table 9. Western blot assay was conducted using standard techniques, with the following primary antibodies; CFIm25 (10322-1-AP, Proteintech or sc-81109, Santa Cruz), REX1 (MA5-38664, Thermo Scientific), CFIm68 (A301-358A, Bethyl), CFIm59 (A301-359A, Bethyl), GAPDH (sc-32233, Santa Cruz), Flag (HT201-01, TRANS), His (HT501-01, TRANS).

## Accession numbers

All the deep sequencing data have been deposited to GEO database with the accession no.GSE178194

## Acknowledgments

This work is supported by a National Natural Science Foundation of China (31970613) and Guangzhou Municipal Science and Technology Project (201803040017) to C. Y.

---

**Figure 1-figure supplement 1.**
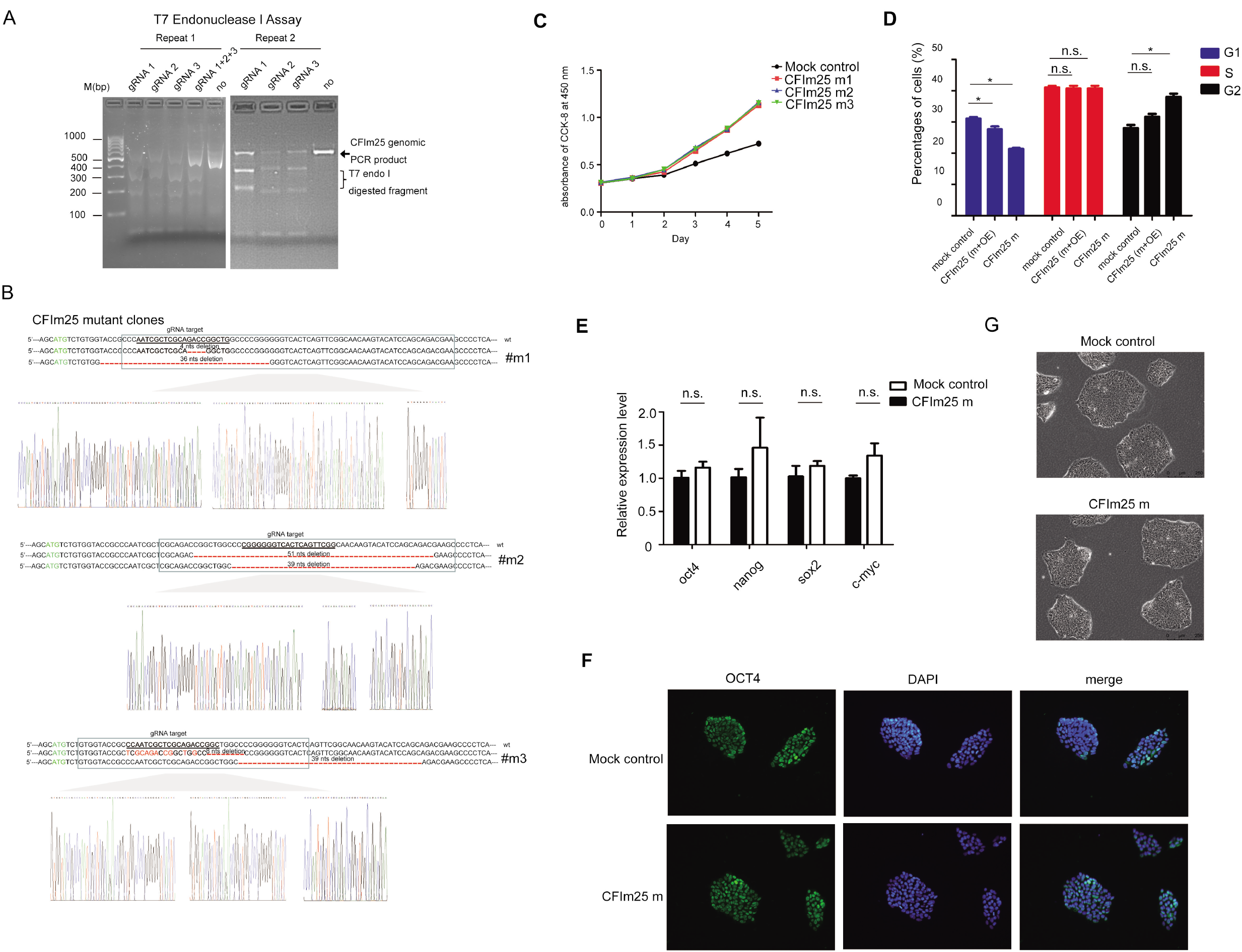
(A) SYBR Green staining of PCR product and DNA fragments resulted from T7 Endonuclease I assay. PCR product amplified from CFIm25 gene locus is indicated by the arrow and digested DNA fragments is indicated by the bracket. Two representative replicates experiments are shown. (B) Sanger sequencing of CFIm25 gene locus to confirm the genomic mutations/deletions in three CRISPR-Cas9 system-mediated H9 cell clones. Start codon ‘ATG’ is colored green, nucleotides colored in red represent mutations, and symbol “-” stands for nucleotide deletion at the corresponding position. Genomic positions targeted by gRNAs are underlined. (C) Cell proliferation rate measurement by CCK-8 kit in mock and three CFIm25-mutant H9 cell lines. The starting cell density in this experiment is 7500 cell per well of 96 well plates. Three independent experiments have been carried out and representative results are shown. (D) Quantifications of the percentages of cells at different stages during cell cycles. The results are from three independent experiments. A representative result is shown in Figure 1D (m: mutant; OE: overexpression). Student’s t-test was used to estimate the significance of the change. *P<0.05; n.s.: non-significant. (E) RT-qPCR analysis of the expression level of four pluripotency-associated markers in mock and CFIm25-m hESCs. (F) Immunostaining analysis of pluripotency marker OCT4 in mock and CFIm25-m hESCs. (G) Phase-contrast images of mock and CFIm25-m hESC clones.

**Figure 2-figure supplement 1.**
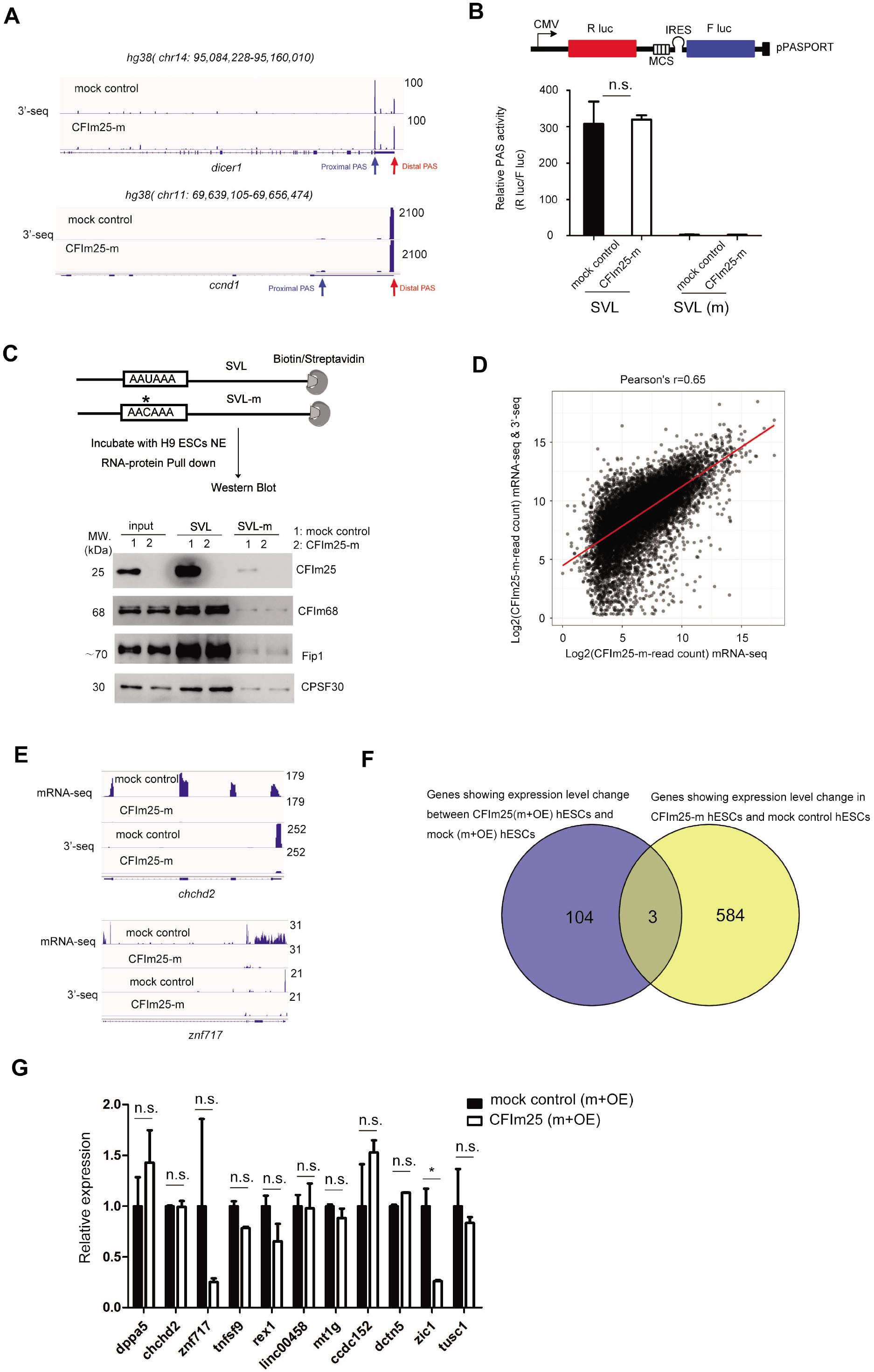
(A) IGV track screen shots showing the 3’-seq results for dicer1 and ccnd1 genes in mock and CFIm25-m H9 cells. Two predominant PASs within 3’UTR regions are indicated with arrows. Proximal or distal PAS are named according to their positions relative to gene 5’ end. (B) Measurement of canonical SVL PAS processing efficiency using pPASPORT system in mock and CFIm25-m cells, SVL-m (AAUAAA core hexamer was replaced as AACAAA) serves as negative control. SVL or SVL(m) PAS were inserted into multiple cloning sites between Renilla luciferase (Rlu) gene and IRES (internal ribosome entry site). Downstream of the IRES is the Firefly luciferase (Flu) gene. Relative PAS processing efficiency was quantified by calculating the Rlu/Flu ratio. Results from three independent experiments are quantified and represented. Student’s t-test was used to estimate the significance: ns: non-significant. (C) Schematic representation of the SVL RNA substrates used in the biotin–streptavidin pull-down assay (top). The AAUAAA hexamer in wild-type RNA substrate and AACAAA in mutant substrate (boxes) are shown. The asterisk is used to highlight the single nucleotide change. Bottom panel shows the western blot results of known core 3’ processing factors in the RNA-biotin based pull-down experiment using nuclear extracts (NEs) prepared from mock and CFIm25-m H9 cells. Two independent experiments have been carried out and representative results are shown. 5% of the lysate was kept as input. The primary antibody for CFIm25 is from santa cruz (sc-81109). (D) Comparison of gene expression profiling by 3’-seq and mRNA-seq. X axis: total read count for each gene in CFIm25-m sample based on mRNA-seq data; Y axis: total read count for each gene in CFIm25-m based on mRNA-seq results of control and 3’-seq analysis. The mRNA-seq read count for a gene in CFIm25-m sample=(mRNA-seq read count for this gene in control H9) x (expression fold change for this gene based on 3’-seq analyses: CFIm25-m/control). Both X axis and Y axis are in log scale. Pearson’s r=0.65. (E) IGV track screen shots showing mRNA-seq and 3’-seq results for chchd2 and znf717 genes in mock and CFIm25-m H9 cells. (F) Venn diagram showing the number of overlapping and non-overlapping genes that display expression level change upon CFIm25 (m+OE) (blue) and CFIm25-m (yellow) (m: mutant; OE: overexpression). (G) Comparison of the expression level of indicated genes in mock (m+OE) and CFIm25 (m+OE) H9 cells. Data comes from RNA-seq analysis listed in Supplemental Table 4. Student’s t-test was used to estimate the significance: *p<0.05; ns: non-significant.

**Figure 3-figure supplement 1.**
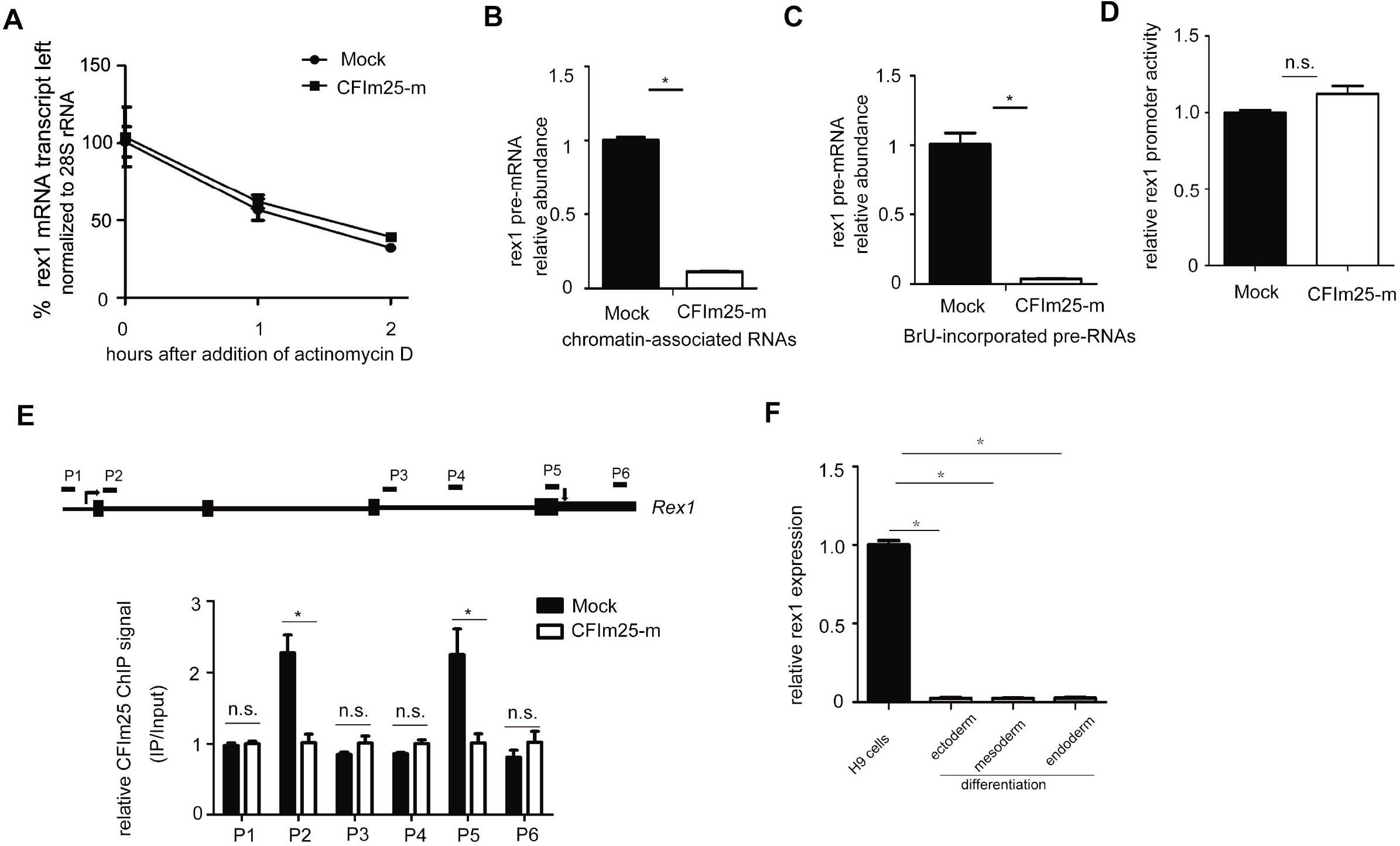
(A) Mock and CFIm25-m (1-3) H9 cells were subjected to Actinomycin D treatment and total RNAs were extracted at the indicated time. RT-qPCR was used to calculate the percentage of the rex1 mRNA left, 28s rRNA serves as normalization control. (B-C) RT-qPCR analysis of rex1 pre-mRNA abundance using chromatin-associated RNAs (B) and BrU-incorporated pre-mRNAs (C) in mock and CFIm25-m H9 cells. gapdh gene product serves as internal normalization control. (D) Comparison of rex1 gene promoter activity in control and CFIm25-m H9 cells using pGL3-basic reporter system. Student’s t-test was used to estimate the significance of the change. ns: non-significant. (E) ChIP-qPCR analysis using primary antibody against CFIm25 and indicated primers targeting different position of rex1 gene locus. For the bar graph, y axis represents the fold change of ChIP signal in mock H9 cells in comparison to that of CFIm25-m cells, x axis stands for the indicated positions across rex1 gene locus. (F) RT-qPCR analysis of rex1 gene expression in undifferentiated H9 cells, and trilineage differentiated cells. Gapdh mRNA was assayed as normalization control. Student’s t-test was used to estimate the significance of the change. *P<0.05.

**Figure 4-figure supplement 1.**
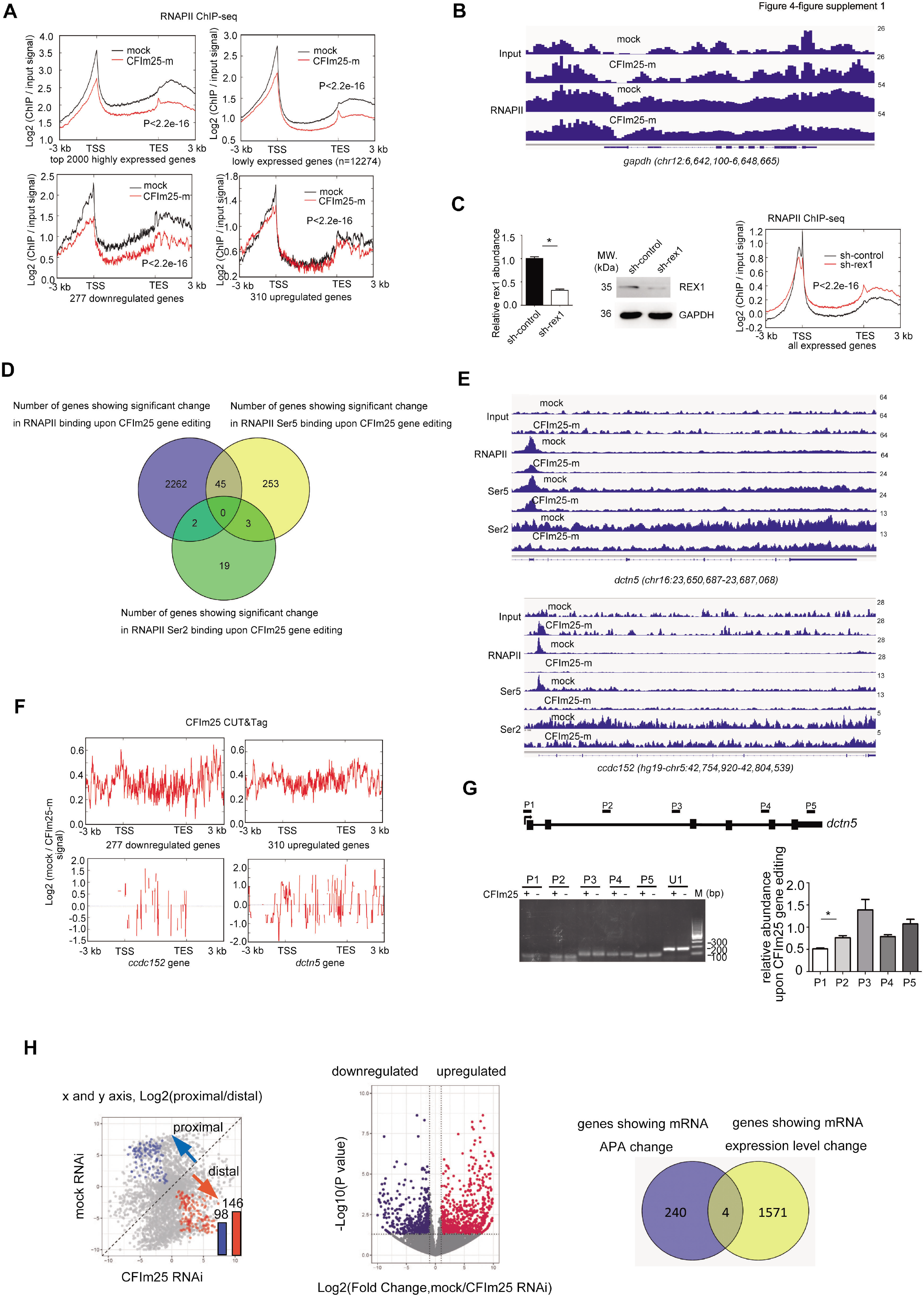
(A) Metagene plots of RNAPII ChIP-seq reads for highly expressed genes (top 2000 genes based on mRNA-seq FPKM value), lowly expressed gene (the rest of the genes), 277 down-regulated genes and 310 up-regulated genes upon CFIm25 gene editing. K-S test was used to examine the significance of the difference between the two plots. (B) IGV track screen shot showing RNAPII ChIP-seq result for gapdh gene in mock and CFIm25-m H9 cells. (C) Metagene plots of RNAPII ChIP-seq reads for actively expressed genes in mock and rex1 RNAi H9 cells. K-S test was used to examine the significance of the difference between the two plots. The rex1 gene knockdown efficiency was estimated by RT-qPCR and western blot analysis. Student’s t-test was used to estimate the significance of the change. *P<0.05. (D) Venn diagram showing the number of genes that displayed differential RNAPII/Ser5/Ser2 binding upon CFIm25 depletion. (E) IGV track screen shots showing RNAPII, RNAPII Ser5 and Ser2 ChIP-seq results for dctn5/ccdc152 gene in mock and CFIm25-m H9 cells. (F) Plots showing the normalized CFIm25 CUT&Tag signals in the specific group of genes (up-regulated or down-regulated genes) or individual gene (ccdcl52, dctn5). (G) Nuclear run-on assay on the nascent dctn5 transcript. The gene structure and the probe positions are indicated on the top. The diagram for the nuclear run-on assay is shown in the Figure 4F. A representative set of RT-PCR data are shown in the middle panel. RT-PCR products from mock and CFIm25-m H9 cells are indicated below each set. ‘CFIm25-’ represents CFIm25-m cell nuclei, whereas ‘CFIm25 +’ represents mock cell nuclei. Bar graph represents RT-qPCR data from three independent experiments. U1 snRNA was assayed as normalization control. Student’s t-test was used to estimate the significance of the change. *P<0.05. (H) Comparison of global mRNA APA (left) and gene expression profiles (middle) in control and CFIm25 RNAi cells using the previously reported dataset. The plots are similar to that in Figure 2A-B. Venn diagram shows that the two groups of genes showing changes do not overlap extensively.

**Figure 5-figure supplement 1.**
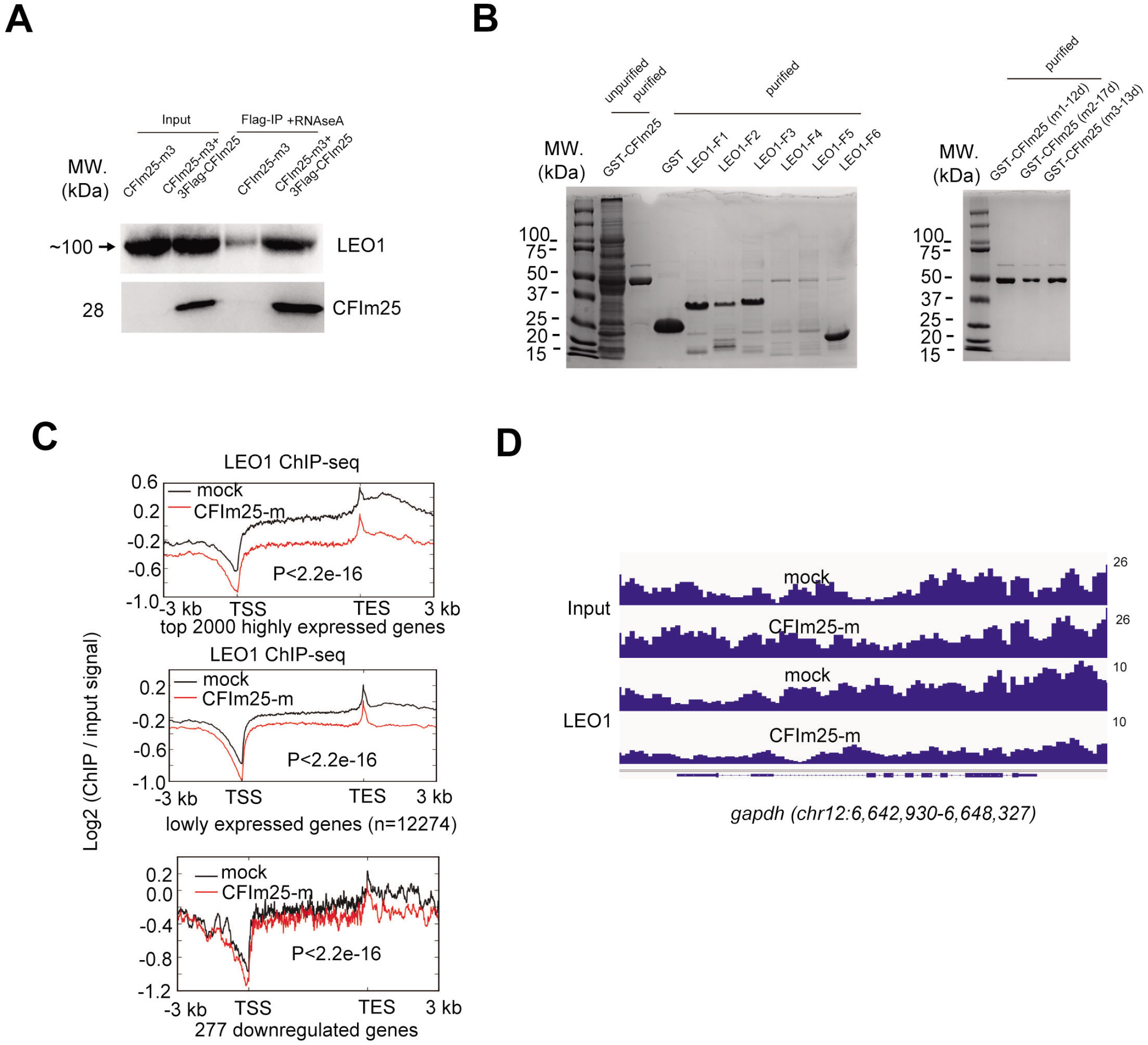
(A) Western blot analysis of the abundance of LEO1 protein in Flag-IPed sample. Flag-IP was performed using extracts from CFIm25-m3 and CFIm25 (m3+3XFIag-CFIm25 overexpression) H9 cells in the presence of 5 ug/ml RNAse A. Input: 1% of the lysates for IP. (B) Commassie blue staining of purified GST-CFIm25, His-LEO1 (truncation fragments 1-6) fusion proteins (left), and three GST-CFIm25 mutants (the N terminus mutations are based on sequences listed in Figure 1-figure supplement 1B). (C) Metagene plots of LEO1 ChIP-seq reads for highly expressed genes (top 2000 genes based on mRNA-seq FPKM value), lowly expressed gene (the rest of the genes), and 277 down-regulated genes upon CFIm25 gene editing. K-S test was used to examine the significance of the difference between the two plots. (D) IGV track screen shot showing LEO1 ChIP-seq result for gapdh gene in mock and CFIm25-m H9 cells.

**Figure 6-figure supplement 1.**
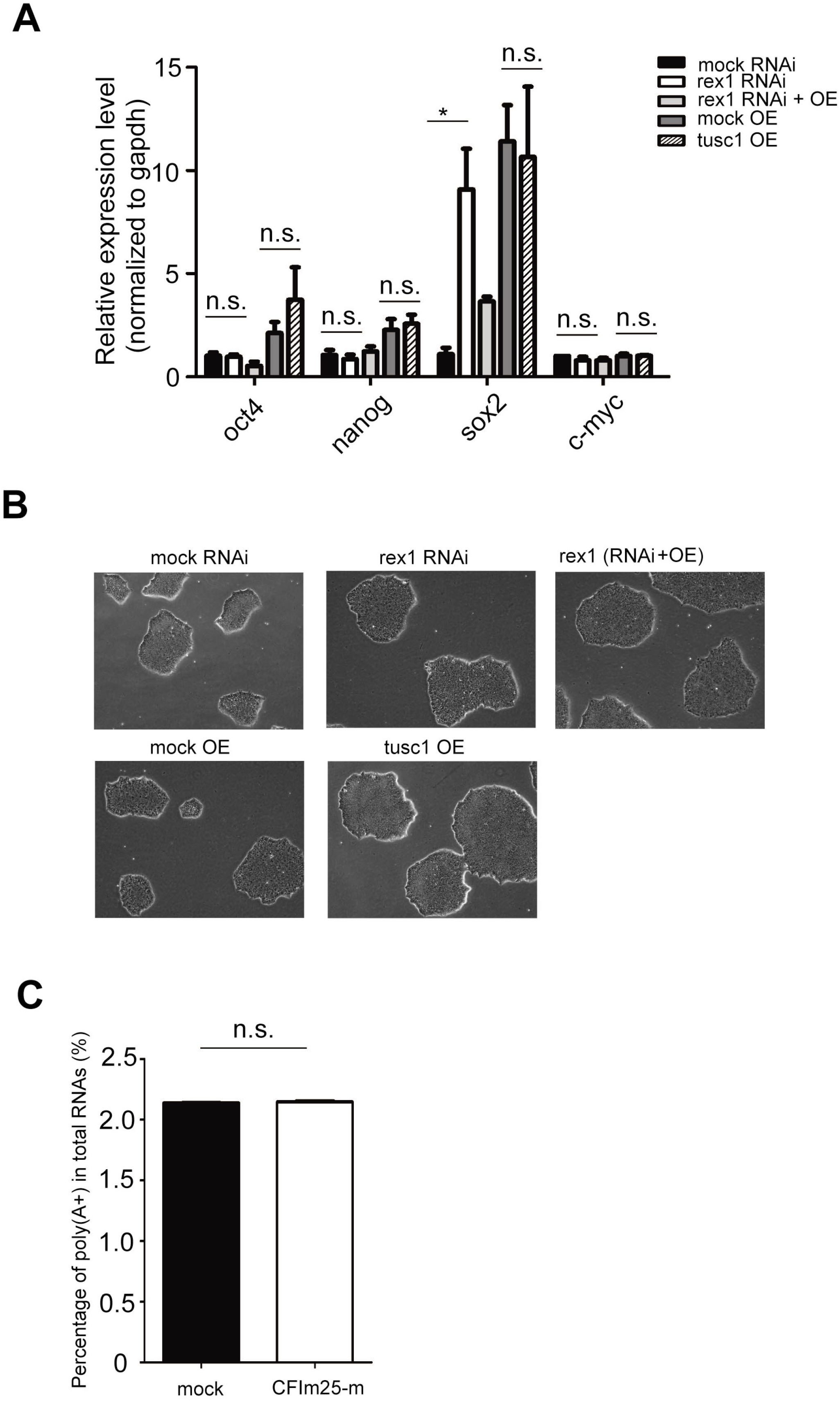
(A) RT-qPCR analysis of the expression level of four pluripotency-associated markers in indicated cell lines. Student’s t-test was used to estimate the significance of the change. *P<0.05; ns: non-significant. (B) Representative phase-contrast images of indicated cell lines. (C) Bar graph showing the percentages of poly(A+) RNAs among total RNAs in mock and CFIm25-m H9 cells. Poly (A+) RNAs were purified by OligodT magnetic beads from total RNAs. Quantification was performed with three independent experiments. Student’s t-test was used to estimate the significance of the change. ns: non-significant.

